# Immune Cell Migration Models Synergize Nuclear Piston, Uropod, and Microenvironment into Hydraulic Cell Engine

**DOI:** 10.1101/2025.09.02.673867

**Authors:** Sami Alawadhi, David M. Rutkowski, Yerbol Tagay, Alexander X. Cartagena-Rivera, Alexander S. Zhovmer, Denis Tsygankov, Dimitrios Vavylonis, Erdem D. Tabdanov

## Abstract

The nucleus and uropod are the largest and most mechanically distinct structures in migrating amoeboid lymphocytes, including NK, B, and T cells. The biophysical properties of these structures may shape the ability of immune cells to navigate dense tissue microenvironments during immune surveillance. Using bead-spring and agent-based cell models, we explore the biomechanical contributions of the nucleus, uropod, septin-templated cortical rings, actomyosin cytoskeleton, and extracellular matrix obstacles to lymphocyte migration. Our results support a migration model in which, following cell-matrix collisions, septins mediate the formation of cortical rings that hydraulically seal cytoplasmic compartments on each side of the passing nucleus, generating a pressure difference that propels the nucleus forward. This hydraulically driven nuclear piston actively enhances migration through confined spaces. Concurrently, the uropod emerging from the peristaltic collapse of rear compartments stabilizes directional persistence and prevents T cell repolarization. We show that such polarity stabilization boosts immune surveillance efficiency. Together, these models redefine the nucleus as an active component of the migratory engine and the uropod as a locomotion stabilizer. Furthermore, the models offer a predictive framework towards engineering of immune cell motility in complex tissue microenvironments with broad implications for cancer immunotherapy, aging, and regenerative medicine.

## INTRODUCTION

Amoeboid migration is a rapid ^1–5^ and energy-efficient ^6–8^ mode of immune and cancer cell migration, operating *via* weak transient ‘focal’ adhesions ^9^ or completely independently of adhesion ^1,10–12^ and proteolysis of extracellular matrix (ECM) ^10,13^. Amoeboid migration is essential for immune surveillance ^14^, but also exploited by various cancers during metastatic dissemination ^14–17^. Therefore, elucidating biophysical, biochemical, mechanobiological, signaling, metabolic, and integrative aspects of amoeboid motility mechanisms is of broad significance across cell biology, immunology, and cancer therapeutics. However, despite its importance, a fundamental mechanistic understanding of amoeboid migration, particularly three-dimensional (3D) migration within physiologically relevant environments, is lacking ^18^.

At its core, amoeboid migration relies on continuous and dynamic changes in cell shape, enabling cells to probe, adapt to, and maneuver through their surrounding microenvironment ^3,12,18^. This morphological plasticity enables amoeboid cells to propel themselves using minimal adhesion, instead leveraging steric, non-adhesive interactions with the ECM as mechanical fulcrums for movement ^10,19,20^. Rather than relying on a singular propulsion mechanism, current evidence derived from studies of immune cells suggests that amoeboid migration encompasses a spectrum of interrelated mechanobiological strategies. The reported mechanisms include the osmotic engine ^21^, paddling ^22^, worrying ^19^, peristaltic ^20^, blebbing ^19,23–25^, and advected cortex percolation ^26^, highlighting the multifaceted and adaptive nature of amoeboid motility across diverse tissue environments.

The extreme plasticity of immune cell migration ^16,27,28^ suggests that lymphocytes have evolved the ability to switch between distinct modes of amoeboid motility in response to changing conditions. This versatility is essential for adapting to the wide range of structural and mechanical environments encountered across tissues and organs during immune surveillance ^16,20,29–31^. Thus, a higher-order comprehensive model of immune cell migration is needed to account for its ability to dynamically balance and/or combine the different mechanisms of amoeboid and amoeboid-like motility, which includes the dynamic balance between cell adhesiveness, contractility, and protrusive activity, collectively defining various forms of amoeboid and amoeboid-like migration^2^.

Historically, the mechanisms of amoeboid migration in mammals have been studied using predominantly dendritic cells (DC) ^1,32,33^, to a lesser extent, T cells ^20,34^, and melanoma cells for cancer research ^19,35^. DCs are considered ‘professional migrators’ ^1,36,37^ due to their highly adaptable mechanobiological properties, which enable extreme elongation, branching, and efficient navigation through confined environments such as dense ECM and endothelial junctions during diapedesis ^37,38^. The exceptional compliance and deformation tolerance of DCs enable them to migrate through tight interstitial spaces with minimal mechanical resistance ^37^ and traction ^39^. While DC migration has been well characterized descriptively and correlatively, the underlying mechanistic (*e*.*g*., biophysical) principles driving their amoeboid locomotion remain poorly understood. Current evidence indicates that DCs migrate using a myosin-dependent, adhesion-independent mode ^1,32,36,40^, tightly regulated by microtubule-mediated spatiotemporal coordination of actomyosin contractility through the microtubules→GEF-H1→RhoA signaling axis ^32^. These facts underscore the need for further mechanistic dissection of amoeboid motility to refine our understanding of immune cells’ migration and surveillance.

Similar to DCs, T cells also rely on the microtubules→GEF-H1→RhoA signaling axis to coordinate their migration ^2^. However, T cells exhibit a distinct motility phenotype characterized by reduced deformability and a compact (rounded or segmented) morphology ^20,34^.This structural profile reflects a different migration strategy: unlike dendritic cells that rely on broad plasticity, T cells adopt a peristaltic treadmilling mode of amoeboid migration. This mechanism is driven by differential actomyosin contractility between segmented cortical compartments, enabling coordinated cytoplasmic flow and forward propulsion ^20^. Peristaltic treadmilling is thought to be particularly effective in environments that do not tightly (physically) confine the cells but rather obstruct cell migration with densely populated discrete obstacles, such as the reticular collagen fibers of lymph nodes or the spleen, where T cells must rapidly move around and probe for antigens presented by dendritic cells ^41^.

In three-dimensional reticular collagen networks, the crowding architecture of the extracellular matrix induces mechanical indentations in the T cell cortex, triggering the localized formation of septin-templated cortical rings. These rings, condensed from filamentous actin and cross-linked with actinin, compartmentalize the cortex into discrete, independently contracting segments ^20^. Such structural segmentation allows T cells to dynamically redistribute cortical tension and shape, adapting to extracellular constraints. As migration proceeds, the cell contracts the rear to form the uropod while simultaneously extending the front compartment, propelling the cell forward in a wave-like motion. This peristaltic cycling of protrusion and retraction enables the efficient translocation of cytoplasm and the nucleus through complex extracellular landscapes, allowing T cells to circumnavigate, evade, and optimize their migration path through dense yet discretely obstructed environments with high efficiency ^20^.

Another key aspect of immune cell migration involves the mechanical roles of the nucleus and the uropod, two structurally dominant and mechanically distinct features of migrating lymphocytes. The nucleus, due to its substantial size and stiffness, poses a significant physical barrier to movement through the physically confining intercellular spaces of solid tissues and vascular walls ^42^. Experimental evidence shows that cells with more deformable nuclei migrate more efficiently under such confinement, whereas nuclear stiffening restricts cell motility ^42–45^. The uropod, a narrow, cylindrical extension at the rear of the cell, is also crucial for immune cell migration, contributing to locomotion, cell-cell interactions, and transendothelial trafficking during immune surveillance ^46^. Recent reports suggest that the uropod acts as a hydrodynamic wind vane, generating stirring effects to aid navigation in fluid environments ^47^. However, the direct contributions of the nucleus and uropod to the complex and multifaceted mechanisms of amoeboid motility remain unclear, highlighting a need for further investigation.

A growing body of evidence suggests that the nucleus functions as a mechanical “piston” ^48–50^, an ECM-pushing “ram” ^48,51–53^, and a “geometric ruler” that senses environmental confinement ^54^. These roles converge into an integrated “piston-ram-ruler” signaling loop, where nuclear deformation provides critical mechanosensory feedback to guide cells through tight spaces ^55^. In the “ram” model, forward thrust is generated by pressure exerted by the nucleus on the surrounding cytoskeleton and plasma membrane, enabling cells to overcome physical barriers and narrow pores ^56^. This concept relies on a mechanical coupling between the nucleus and the cytoskeleton, anchored *via* adhesion to the external fulcrum, *i*.*e*., the extracellular matrix (ECM), which translates cytoskeletal traction into nuclear propulsion. Specifically, this coupling is mediated by nuclear adaptor complexes such as SUN1/2 and Nesprin-1/2 ^57^, the LINC complex ^58,59^, and additional linkages including RanBP2-BicD2 and Nup133-CENP-F ^60–62^. Cytoskeletal forces may be generated by either actomyosin contractility ^63,64^ or microtubule motors ^61,65^.

Several models posit that cytoskeletal forces that drive the nucleus’s forward thrust are converted into hydrostatic pressure at the nucleus’s front, facilitating lobopodial inflation and cytoplasmic flow in an amoeboid manner ^50,63,64^. Alternatively, hydrostatic or solid-state pressures from the rear, generated by compression of trailing microtubules, may also push the nucleus forward ^53^. Regardless of their mechanistic variations, the listed models converge on a common principle: the nucleus acts as a mechanical element of the machinery driving 3D cell migration in structurally dense environments.

The described classical framework, however, presents a *paradox* for immune cells. Unlike mesenchymal cells, T lymphocytes lack stable actomyosin stress fibers ^66^ and robust ECM adhesions ^12,67^, both essential for traction-based nuclear propulsion. Instead, the T cell cortex features a dynamic, loosely organized actomyosin network. This structure cannot be effectively and directly coupled to the nucleus to apply traction, but instead allows the nucleus to slide rapidly along the cortex during migration through reticular collagen networks ^20^. This observation challenges the applicability of cortex-to-nucleus adaptor-mediated force-transmission models in T cells. The absence of robust cytoskeleton-ECM linkages suggests that T cells may rely on internally generated pressure gradients rather than traction-based mechanisms. If true, this *points to an alternative*, currently undescribed, nuclear-adaptor-independent mechanism for transmitting cytoskeletal forces to the nucleus.

Notably, studies with DCs indicate that these cells form rear-end uropod-like compartments and rely on myosin accumulation to propel the nucleus through constrictions in microfluidic devices. This process is driven by actomyosin-mediated cortical forces, which generate hydrostatic pressure to push the nucleus forward ^59,68,69^. However, the emerging possibility that nuclear movement can be primarily driven by hydrostatic pressure difference between the front and rear cytoplasmic compartments, physically separated from each other by the nucleus, remains largely unexplored.

There is sufficient experimental evidence supporting the idea that the tail-like extension of the rear compartment of an immune cell, *i*.*e*., the uropod, may have a special role in migration. For example, uropod provides polarized localization of contractile non-muscle myosin II ^70^ to generate retraction forces at the cell rear ^71–74^, as well as clusters of ICAMs, CD44, and PSGL-1 to facilitate transient cell-ECM and cell-cell adhesions ^46^. Furthermore, the direct labeling of the mechanically tensed F-actin in cytotoxic T cells with F-actin conformation probes based on the CH domain of utrophin clearly outlines uropod as the most tension-loaded part of the cell cortex ^75^. Additionally, the uropod often hosts a microtubule-organizing center (MTOC) ^46,76^ to reinforce cell polarization ^77^. The myosin polarity in lymphocytes comes from the opposing activity distributions of RhoA (at the uropod) and Rac1 and Cdc42 (at the cell front) ^78^, allowing for increased F-actin polymerization at the cell’s expanding front ^79^. Consequently, as shown in DCs, the uropod serves as the final destination of retrograde cortical actin flow, which is driven by myosin II contractility that compacts the cortical cytoskeleton ^33^. Notably, these uropod-related polarization mechanisms, observed in relatively small immune cells, can also be recapitulated in immune cell-sized fragments of larger, non-immune cells ^26^. These observations suggest the existence of a fundamental, scale-dependent governing principle that links cytoplasmic hydrostatic forces and the structural and mechanical dynamics of the contractile cortex, ultimately facilitating a stable amoeboid polarization of small lymphocytes.

This computational study integrates the key governing principles of nucleus-cortex interactions and the uropod function in migrating amoeboid cells to develop a comprehensive, higher-order model of immune cell migration. A predictive mechanobiological model of immune cell migration can guide the re-engineering approaches to enhance T- and NK cell motility by tailoring their migratory properties for the specific tasks of immunotherapies. Such a model can also help to investigate the interplay between the intracellular structural and mechanical properties of amoeboid immune cells and the mechano-structural complexity of their surrounding microenvironment in both healthy and cancerous tissues. By incorporating insights from the cytoskeletal organization, actomyosin contractility, and nucleus deformability, we provide a mechanistic framework for capturing the adaptive strategies that amoeboid cells employ to navigate through confining and heterogeneous environments.

## RESULTS

### Basis of the T cell amoeboid motility model

Septin-templated cortical rings form in T cells as a cytoskeletal cortex response to surface indentations at cell-obstacle collision sites ^20^. Such cortical conformity to the 3D environment induces T cell segmentation, which enables amoeboid peristalsis, propulsion, and circumnavigation in confined or crowded spaces. Cortical rings can also engage the translocating nucleus to produce a tight seal around it during passage through restricting obstacles **(Figure 1A)**. Such a seal suggests a conversion of the nucleus into a hydraulic piston that can be driven by pressure gradients between collapsing rear and expanding front cellular segments **(Figure 1B-1)**.

**Figure 1.**
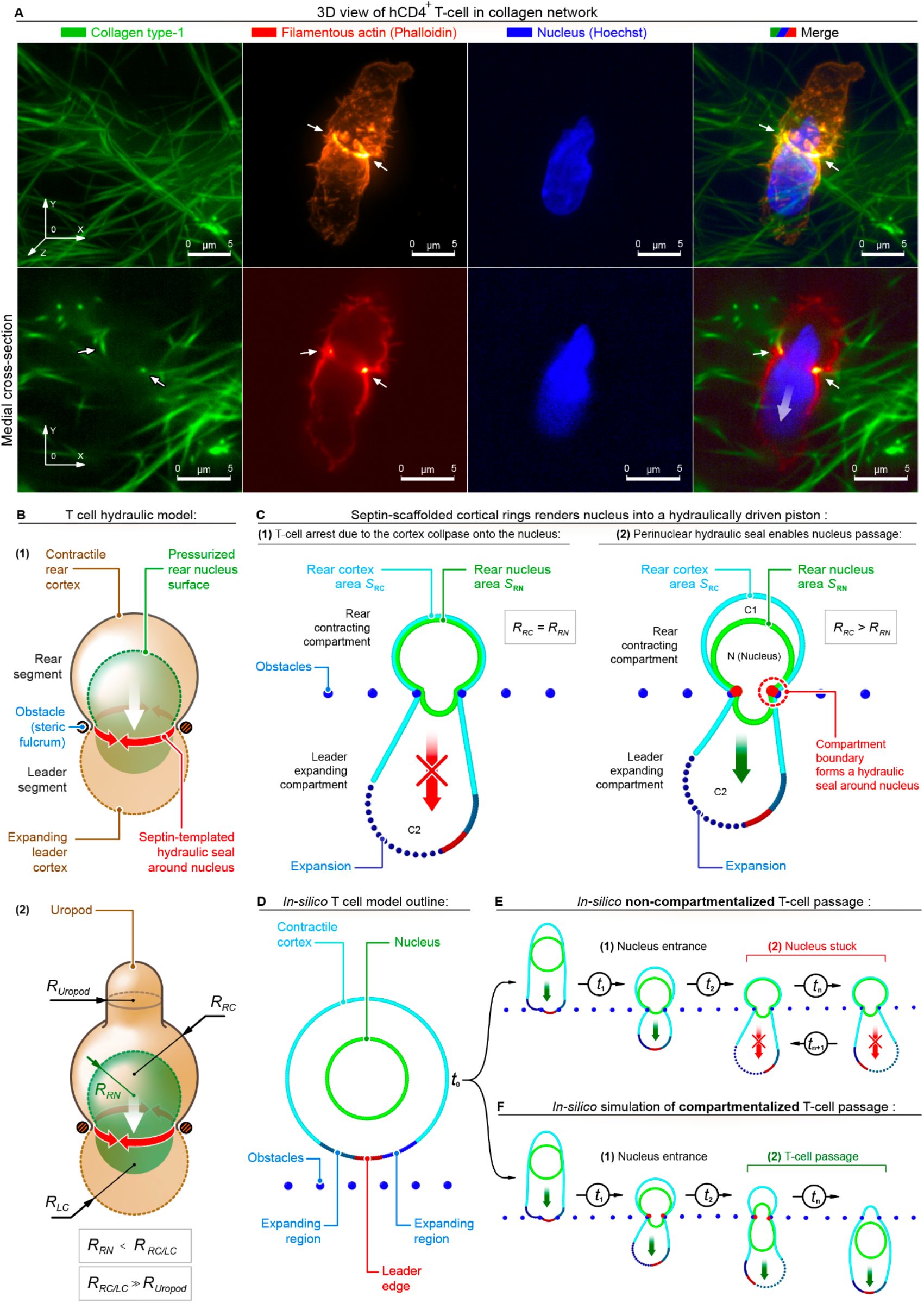
*In-silico* model of hydraulic enhancement of nucleus propulsion in T cells navigating obstacles. **A** - (*Top*) maximum projection of 3D reconstruction and (*Bottom*) cross-section views of the migrating amoeboid human CD4^+^ T cell within a reticular collagen type-1 network. *Arrows denote the collagen obstacles and the obstacle-induced cortical ring*. **B** - T cell hydraulic model: (**1**) Schematic representation of the simplified two-chamber hydraulic model of a T cell. An obstacle-induced cortical ring, templated by septins, acts in concert with the passing nucleus to form a sealed, hydraulically driven piston system. This system divides the cytoplasm and cortex into distinct front and rear compartments, with the rear compartment generating higher hydrostatic pressure than the leading compartment. The resulting pressure differential drives forward translocation of the nucleus as a hydraulic piston. (**2**) A more detailed hydraulic T cell model incorporating the uropod. The compact, tail-like uropod extends from the rear compartment and is characterized by a small principal radius (*R*_*Uropod*_), compared to the rear (*R*_*RC*_), and leader (*R*_LC_) compartments. Due to the uropod’s inherently dynamic compaction into a small-radius, high-contractility appendage, driven by its elevated actomyosin activity, according to Laplace’s law, its cortex can actively induce an elevated hydrostatic pressure in the rear compartment. This configuration may allow the uropod to sustain high hydrostatic pressure behind the nucleus, thereby facilitating forward propulsion of the nuclear piston during amoeboid T-cell migration. **C** - Comparative computational models of T cells without (**1**) and with (**2**) compartmentalization highlight the effects of hydraulic compartmentalization during obstacle circumnavigation. In both cases, the model stipulates a higher hydrostatic pressure in the rear compartment (the uropod is not explicitly modeled). **D** - 2D model of T cell before obstacle encounter: The contractile cortex is polarized with a designated leading region capable of expansion (bleb formation). The nucleus is modeled as an elastic and deformable object with a nearly constant area (incompressible volume). The cytoplasm maintains a nearly constant area, representing an incompressible volume. **E** - Non-compartmentalized T cell model: T cells without compartmentalization fail to navigate through narrow passages between obstacles because forces from the rear contractile cortex are depleted at the immediate proximity to the rear nucleus, leading to an arrest in migration. **F** - Compartmentalized T cell model: Successfully navigates obstacles by employing hydraulic force transmission from the rear contractile cortex to the nucleus, facilitating movement through constricted spaces.

The nucleus-driving hydrostatic pressure gradient across the T cell can arise from the diffusion ^80,81^ and mechanical ^20^ separation of the rear and leading compartments by the septin-templated cortical ring. In this case, a difference in cortical contractility can be directly attributed to the sporadically established difference in myosin activity between the segments ^20^. For a stable contractility gradient T cells can utilize structural adaptations, such as the uropod, a narrow (2-3 µm-wide) tail-like structure at the cell rear **(Figure 1B-2)**. Uropod features a substantially smaller curvature radius (*R*_*Uropod*_) than the adjacent rear compartment (*R*_*RC*_) ^47,82^ and often exhibits elevated actomyosin activity (RhoA accumulation) ^70^ and cortical tension (tensed F-actin molecular probe recruitment) ^75^. According to Laplace’s law, this geometry and tension can amplify hydrostatic pressure in the rear compartment **(Figure 1B-2)**. The uropod thus can act as a pressurizing unit, maintaining a pressure differential that drives forward nuclear translocation, which will be explored in detail in the next sections. Although our following computational cell model does not *explicitly* include the uropod as a part of the rear compartment, its effect is accounted in the model as a robust pressure gradient between the front and rear compartments upon the encounter of the nucleus with the septin-mediated cortical ring at the constriction and, thus, capturing the efficient nuclear piston-driven motility during amoeboid T cell migration.

### Compartmentalization facilitates the migration of T cells

First, we examine the outlined hypothesis that the septin-templated cortical rings, acting as the perinuclear seals that separate the compartments, can assist in propelling the nucleus as a hydraulic piston during amoeboid T cell migration. For that we compare the simulated two-dimensional (2D) migration of T cells through the obstacles in the absence **(Figure 1C-1, Movie 1)** and in the presence **(Figure 1C-2, Movie 1)** of a hydraulic seal around the passing nucleus. The hydraulic seal is represented by two-point boundaries on either side of the cell at the sites of cell-obstacle interactions. A full description of this model can be found in the **Methods** section, while the main assumptions and resulting cell behaviors are outlined in the following paragraphs.

Simulations of a T cell encountering and passing between the obstacles spaced at half the nuclear diameter, and without a perinuclear hydraulic seal, display an escape (*i*.*e*., leakage) of the cytoplasm from the rear segment into the leader segment. The leakage causes the rear cortical compartment to collapse directly onto the nucleus **(Figure 1C-1)**. Since the contractile, *i*.*e*., phosphorylated form of myosin is absent from cortical regions adjacent to the nucleus ^20^, we assume such collapse abolishes contractile forces in the rear segment. As a result, the T cell becomes stuck in the narrow passage between the simulated obstacles (*e*.*g*., collagen fibers) due to the low deformability of the nucleus and the lack of myosin-driven contractility in the cortex that collapses onto the nucleus.

In contrast, simulations of T cell motility that include the formation of a hydraulic seal near obstacles and around the nucleus demonstrate successful migration **(Figure 1C-2)**. In this case, the pre-existing incompressible volume of the cytoplasm in the rear contracting segment is depicted in the 2D model as a nearly constant area C1 **(Figure 1C-2)**. The preserved cytoplasm volume enables the rear cortex to project contractile pressure directly onto the posterior nuclear surface, generating a hydraulic “booster” effect. The resulting hydraulic “booster” effect, in coordination with the expanding front segment, facilitates effective nuclear translocation and forward propulsion through the confined space.

The main assumptions of the simulation are shown in **Figures 1D** and **1E**. The model distinguishes between a contractile rear cortical region and a cortical leader region at the cell front. The leading region of the advancing T cell is flanked on either side by ‘blebbing’. The expandable regions are parts of the cortex that can stretch and expand into a new growing segment under hydrostatic pressure **(Figure 1D)**. The nucleus is also shown and is stipulated to feature a higher mechanical rigidity and incompressible volume (nearly constant area in 2D). Its centering is maintained by weak spring forces, which emulate the stabilizing influence of microtubules and intracellular organelles ^83,84^.

Figure 2 further illustrates the proposed role of the septin-templated seal in forming a hydraulic piston, as well as its computational representation. Septin-templated cortical rings **(Figure 2A)** form at sites of cell surface indentation created by surrounding obstacles **(Figure 2B-1)**, in effect producing a stabilized circular opening for the passing nucleus **(Figure 2B-2)**, regardless of the geometry of the confining environment. In addition to preventing excessive nuclear deformation, this geometric constraint facilitates the formation of a tight cortical seal around the nucleus. In the simulations of T cell transmigration, this seal enables functional compartmentalization of the cytoskeleton and a pressure difference between the rear and front segments. Specifically, the presence of the seal results in the mechanical isolation of the rear contractile cortex, which generates elevated pressure on the posterior side of the nucleus compared to the expanding front segment **(Figure 2C**, Movie 1).

In contrast, removal of the seal results in pressure equilibration across the cell due to unrestricted cytoplasmic flow, leading to a uniform pressure distribution around the nucleus. The loss of hydraulic pressure on the nucleus ultimately allows the collapse of the rear cortex directly onto the nucleus, disrupting rear-segment contractility and impairing forward propulsion **(Figure 2D, Movie 1)**.

**Figure 2.**
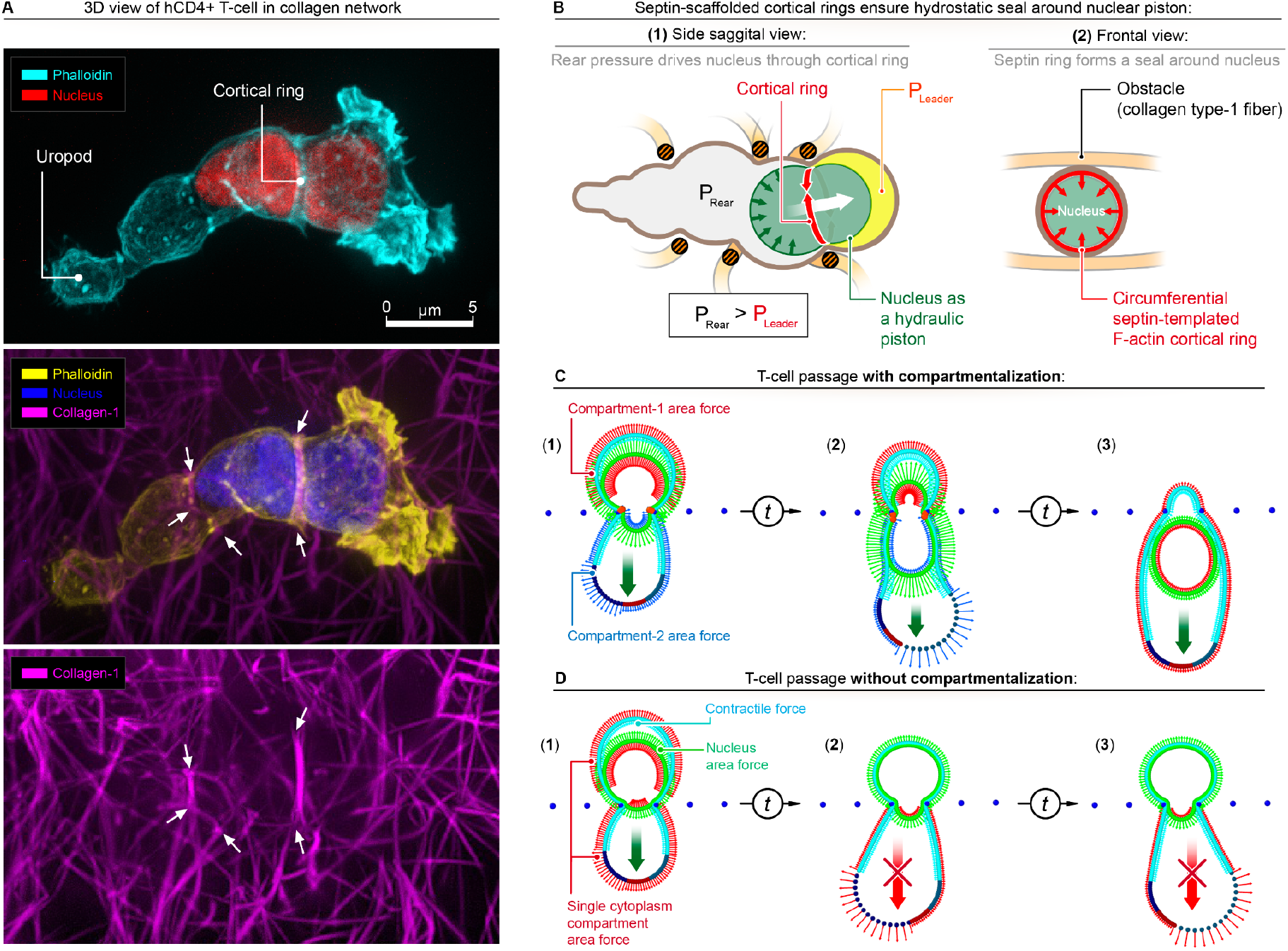
Analysis of hydraulic force transmission between the rear contractile cortex, nucleus, and expanding leading segment. **A** - 3D reconstruction of the migrating amoeboid human CD4^+^ T cell in reticular collagen type-1 network. *Arrows denote obstacles and obstacle-induced cortical rings*. **B** - (**1**) A Schematic of the T cell hydraulic model highlights the hydraulic cortical seal around the nucleus, which forms as a septin-mediated response to the obstacle-induced cortical indentation. (**2**) The cortical ring encircles the nucleus, separating the migrating amoeboid T cell into two hydraulically isolated compartments. The difference in the hydrostatic pressure on either side of the nucleus within their corresponding compartments renders the nucleus into a hydraulically driven piston. **C** - With compartmentalization, the hydraulic force transmission from the rear segments’ contractile force to the rear nucleus surface enables effective nucleus piston propulsion through the spatially constraining obstacles. **D** - Without compartmentalization, cytoplasm escapes from the contractile rear segment into the leader compartment through the available lumens between the cortex and the nucleus without pushing the latter forward. As a result, the rear cortical segment collapses onto the nucleus’s rear surface, only stretching around the nucleus without hydraulic transmission of contractile forces. The lack or low efficiency of cortical force utilization leads to the nucleus being stuck at the entry of the confining obstacles.

In summary, the hydraulic seal surrounding the passing nucleus amplifies the transmission of contractile forces from the rear segment to the posterior surface of the nucleus. Because the contractile cortex has a larger surface area than the nucleus, this configuration enables the hydraulic-based transmission of contractile forces onto the nucleus in the direction of the constriction, effectively enhancing the nucleus’s ability to propel forward through spatially confining obstacles **(Figure 2C)**. In the absence of such compartmentalization, the net cytoplasmic force is not directed along the nuclear passage line. Instead, we assume it to be negligible as the cytoplasm escapes from the rear segment into the leader segment through the gap between the cell cortex and the nucleus.

This cytoplasmic flow results in the loss of the pressure gradients between segments and in the collapse of the rear cortex directly onto the nucleus. In such cases, the cortex in an unmediated physical contact with the nucleus could, at best, directly apply its mechanical force to the nucleus. However, our data, provided below, indicate that such direct force is unlikely because myosin phosphorylation is substantially reduced at the nucleus-cortex interface. Consequently, the nucleus fails to translocate and becomes obstructed by the surrounding constraints **(Figure 2D)**.

### Nuclear positioning defines peristaltic dynamics in amoeboid T cells

To investigate how nucleus localization affects contractile patterning and cell segmentation during amoeboid migration, we analyzed the 3D architecture of human CD4^+^ T cells migrating within dense collagen matrices. We used high-resolution imaging and computational segmentation to examine the density distribution of F-actin, contractile non-muscle myosin II (*i*.*e*., Ser 19 phosphorylated light chain of non-muscle myosin II), and nucleus **(Figure 3A-1)**, revealing the structural layout of non-contractile and contractile cortical compartments relative to the nucleus.

**Figure 3.**
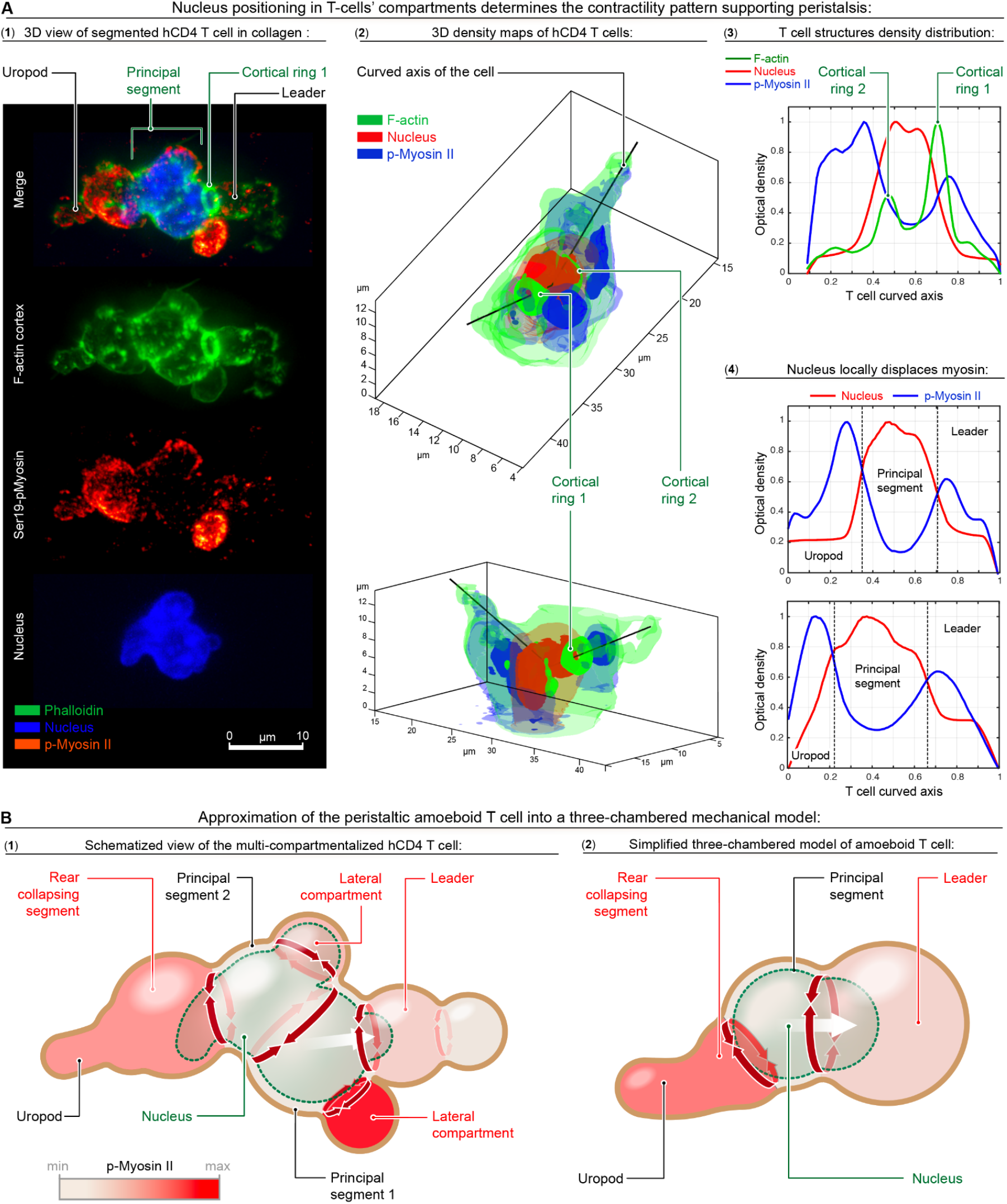
Nuclear positioning in amoeboid T cells regulates compartmentalized peristaltic contractility. **A** - Structural and quantitative analysis of actomyosin organization and nuclear positioning in peristaltically migrating hCD4^+^ T cells embedded in 3D collagen: (**1**) – 3D microscopy image of an hCD4^+^ T cell showing cortical F-actin (*green*), nucleus (*blue*), and Ser19- phosphorylated light chain of non-muscle myosin II (p-Myosin II; *red*). Cortical rings subdivide the cell into three morphological and functional compartments: the rear compartment, which includes the uropod, the principal segment, and the leading segment. (**2**) - 3D volumetric reconstructions reveal that the presence of the nucleus in the principal segment spatially excludes p-myosin II, creating a central region of low contractility. (**3**) - Mean optical density profiles along the T cell’s curved axis demonstrate distinct spatial compartmentalization of contractility and nuclear positioning. p-Myosin II is most enriched in the rear segment (with the uropod), followed by a moderate enrichment in the leading segment, with the lowest enrichment observed in the principal segment containing the nucleus. Segment boundaries correspond to F-actin density spikes marking the positions of cortical rings. (**4**) - Comparative density profiles of the nucleus (*red*) and p-Myosin II (*blue*) along the cell axis further illustrate that the nucleus displaces cortical p-Myosin II in the principal segment. Enrichment of p-Myosin II is observed flanking the nucleus, within the rear and leading segments. **B** - Schematic and mechanical models of peristaltic amoeboid motility in hCD4^+^ T cells: (**1**) - Schematic representation of the cell shown in panel A-1, depicting a nucleus-centered, multi-chambered contractile architecture. The principal (nucleus-containing) segment is flanked by contractile peripheral segments (the rear and leading ones). Color intensity reflects the relative concentration of cortical p-Myosin II. (**2**) - Mechanical approximation of the peristaltic architecture as a simplified three-chamber model, illustrating the interrelation between force generation and nuclear positioning. Coordinated cycles of contraction and expansion drive forward migration through confined environments. This integrative structural and mechanical model supports the concept that nuclear positioning governs compartmentalized actomyosin dynamics, enabling effective peristaltic motility of T cells in complex 3D microenvironments.

Computational segmentation using isosurfaces (points with equal intensity values in a 3D imaging data) shows a consistent organization in which the compartments, occupied by the nucleus, and termed *the principal segments*, are devoid of contractile non-muscle myosin II (p-Myosin II). The principal segments routinely separate two distinct p-Myosin II-enriched compartments: a rear compartment with the uropod and a forward-facing leader compartment. Structurally, 3D density maps of F-actin, nucleus, and p-Myosin II along the cell’s curved axis show two distinct cortical rings **(Figures 3A-1, 3A-2**, and **3A-3)**, highlighting the described cell segmentation. The anterior cortical ring-1 appears ahead of the nucleus, while the posterior cortical ring-2 is localized just behind it, outlining the boundaries of the principal, nucleus-containing segments **(Figure 3A-3)**. Notably, the principal segments exhibit a localized depletion of contractility, detected visually as a depletion of p-Myosin II density at the regions with the nucleus along the cell’s curved axis **(Figure 3A-4)**. This spatial segregation suggests that the nucleus passively partitions the contractile machinery, effectively acting as a mechanical barrier that splits the cell body into rear and front compartments. Such an effect could be attributed to the displacement of the large volume of cytoplasm that contains the pool of myosin-regulating molecular signaling machinery by the nucleus, and also to the effects of the nucleus acting as the Ca^2+^-releasing and uptaking structure ^54,85–88^. Thus, inherently, the nucleus-induced depletion of contractile myosin leads to the loss of the cortical contractility within the nucleus-receiving compartment.

The repositioning of the nucleus initiates a peristaltic cycle in the T-cell cortex, driven by spatially segregated differences in myosin contractility. Specifically, the p-Myosin II-enriched rear segment continuously compacts into the uropod ^20^ **(Figures 3A-1** and **3B-1)**, generating a contractile force that propels the nucleus forward. In contrast, the principal (receiving) segment loses contractile p-Myosin and expands in response, enabling nuclear accommodation. As the nucleus progresses further into the newly formed leader segment, we observe a progressive depletion of p-Myosin II in this compartment, with consistently lower levels compared to the uropod **(Figure 3A-4)**. The loss of myosin-driven contractility in the receiving leader segment facilitates its expansion, allowing for continued forward nuclear translocation and becoming the new principal compartment, hence sustaining the peristaltic cycle. Notably, the rear segment comprises both the collapsing cortical compartment and the geometrically compacted uropod **(Figure 3B-1)** ^20^. Such structural density and geometric compaction of the rear segment could impose a high resistance to re-expansion, preserving tension that prevents backflow and reinforces forward propulsion of the nucleus, functioning as a contractile “valve” in the hydraulic piston mechanism that drives T-cell migration.

To formalize these morphological observations, we use a simplified, three-chambered model of the amoeboid T cell **(Figures 3B-1** and **3B-2)**. The nucleus demarcates three functional compartments: (1) a *rear collapsing segment* associated with high p-Myosin II and contraction forces, (2) a *principal segment* housing the nucleus with reduced contractility, and (3) a *leader* compartment that gradually loses actomyosin contractility and expands into an ECM opening as the nucleus enters this compartment. We also observe lateral compartments, which represent the potential alternatives for the leader segments. However, only one segment that receives the nucleus will become the principal compartment. Thus, despite the structural complexity of the peristaltic T cell **(Figure 3B-1)**, it can be simplified into a three-chambered peristaltic system.

Together, the data and model suggest how the nuclear positioning may serve as a spatiotemporal regulator of peristaltic contractility by spatially constraining actomyosin activity within a segmented (differentially pressured) T cell body.

### Nuclear piston enhances T cells’ ability to infiltrate extremely confining environments

So far, we have described how the cytoskeletal forces create hydrostatic pressure and forward thrust on the nucleus. Here, we explore whether the hydraulically driven nuclear piston mechanism influences the ability of T cells to infiltrate deep into regions of high obstacle density. This question is clinically relevant, as the degree of T cell infiltration into densely packed solid tumors ^89,90^ or fibrotic stromal tissues (*e*.*g*., collagen-rich regions) is highly variable and remains poorly understood, despite being recognized as a critical bottleneck limiting the efficacy of immunotherapies ^91,92^. To address this, we applied the computational model depicted in **Figure 1** to a scenario in which T cells migrate through a field of discrete “collagen fibers” arranged with a spatial density gradient, so that obstacle density increases proportionally with distance from the entry point. We simulated T cell migration through this field both with **(Figure 4A, Movie 2)** and without **(Figure 4B, Movie 2)** the formation of a hydraulic seal, enabling an effective nuclear piston. The results demonstrate that hydraulic compartmentalization significantly enhances T cell penetration into high-density regions of the obstacle field, supporting the notion that the nuclear “piston” mechanism is essential in enabling efficient navigation through densely confining environments.

**Figure 4.**
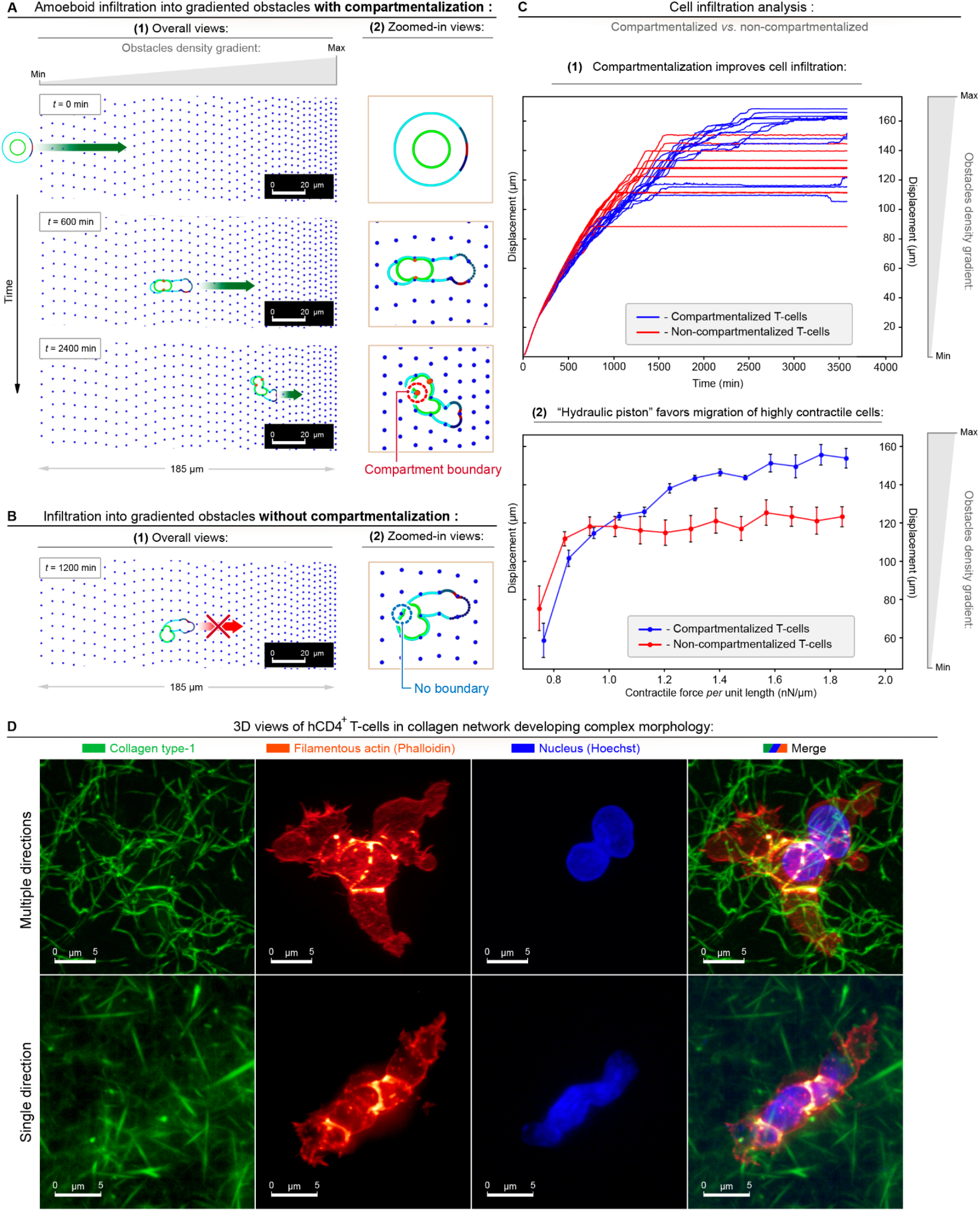
Computational modeling and analysis of the role of the T cell compartmentalization during their migration throughout the progressively increasing obstacle density. **A** - (**1**) Time series of the compartmentalized T cell migration into the increasing obstacle density gradient. (**2**) Enlarged zoomed-in depictions of the modeled T cells at the corresponding time points. **B** - Simulation of the non-compartmentalized T cell displays an early cell migration arrest in the midrange of the obstacle density. **C** - (**1**) Individual T cell penetration curves, *i*.*e*., displacement *vs*. time, into the increasing density of obstacles highlight that most compartmentalized T cells infiltrate into the regions of higher obstacle density. At the same time, the non-compartmentalized cells experience an early arrest within the mid-range of the obstacle density. (**2**) - Compartmentalization of the T cells ensures a more effective accumulation of the tensile cortex energy, enabling a more effective propulsion of the nucleus in dense obstacle environments. With an increase in the obstacle density, the compartmentalization of each T cell also increases, enhancing the engagement of the nuclear “hydraulic piston” that drives cell propulsion deeper into the obstacle density gradient. **D** - Comparative 3D visualization of the hCD4^+^ T cells migrating in dense collagen matrix: *Top* - trident-shaped multi-segment T cell indicates a T cell in the state of exploration of the multiple directions for the amoeboid migration path; *Bottom* - linear multi-segment T cell in a state of unidirectional migration.

Analysis of a large set of simulations of T cell migration within the obstacle gradient reveals a trade-off between unconstrained cytoplasmic flow and nuclear propulsion, governed by hydraulic compartmentalization, *i*.*e*., a hydraulic nuclear piston. The model incorporates the principle of nucleus positioning at the center of the migrating T cell, which is caused by the previously reported weak and steric interactions with microtubules and intracellular organelles ^83,84^ These interactions are modeled as a weak spring force along the direction of the center of mass of the nucleus and cell cortex beads (**Model Methods**). This centering mechanism contributes to the propulsion of the nucleus, causing it to move to the center of the cell during migration in obstacle-free environments. However, centering is insufficient for nuclear propulsion in obstacle-dense regions (**Figure 1E**). In regions of low obstacle density, non-compartmentalized T cells exhibit faster migration due to an efficient rearrangement of the unrestricted cytoplasm and centering of the nucleus within the moving cell **(Figure 4C-1**, non-compartmentalized T cells). However, T cell migration stalls when it encounters areas of higher obstacle density that require substantial nuclear deformation, and the centering forces are insufficient for nuclear passage through these areas. In this situation, compartmentalized T cells, equipped with a hydraulic seal between the cortical rings and the nucleus, utilize the gradient of cortical contractility **(Figure 3)**, migrate more efficiently, steadily, and deeper into high-density regions **(Figure 4C-1**, compartmentalized T cells). These findings highlight the crucial role of nuclear-cortical hydraulic sealing in facilitating effective amoeboid migration through structurally complex and physically constraining environments.

The model suggests that increasing the density of extracellular obstacles around an amoeboid T cell enhances a mechanical synergy between the obstacles, the nucleus, and obstacle-induced hydraulic seals **(Figure 1A)**. Denser environments enhance the formation and number of obstacle-induced septin-templated cortical rings (*i*.*e*., hydraulic seals) that compartmentalize the cytoplasm and prevent the dissipation of contractile force across the cell body. The enhancement of spatial compartmentalization further amplifies the hydraulic piston mechanism that converts cortical contractility into the directed (forward) propulsion of the nucleus.

Modeling results in **Figure 4C-2** illustrate the relationship between the preset T cell cortical contractile force and its migration efficiency, quantified as displacement through a density gradient of physical obstacles, in cells with and without compartmentalization. As shown in **Figure 4C-2**, T cells with low cortical contractile force per length (0.8-1.0 nN/μm) migrate more effectively without compartmentalization, which suggests weak contractile input cannot generate sufficient pressure to drive the nucleus through sealed compartments. In this regime, sealing impairs leader extension and stalls migration. These findings highlight that cortical rings are only advantageous above a critical contractility threshold, beyond which they enable efficient force transmission and directional nuclear translocation through confined environments. Thus, we propose that the formation of septin-templated cortical rings and a hydraulic nuclear piston constitutes a self-regulatory migration mechanism that achieves its full potential only in the presence of high-density obstacles.

In contrast, the absence of a cortical seal permits cytoplasmic flow around the nucleus, allowing the leading compartment’s expansion and advancement, resulting in the overall cell displacement, albeit inefficiently and without pressure-driven nucleus propulsion. This results in a complete cell arrest in more dense obstacles, as the nucleus gets stuck. However, in the higher range of contractile forces (1.0-1.8 nN/μm), compartmentalized T cells display significantly enhanced displacement relative to non-compartmentalized ones. This shift reflects the full engagement of the hydraulic piston mechanism, where elevated cortical contractility generates sufficient pressure within the rear cytoplasmic compartment, sealed by septin-templated cortical rings, to actively drive the nucleus forward. These findings suggest a force threshold above which compartmentalization becomes advantageous, enabling effective translation of actomyosin-generated forces into forward nuclear translocation and overall cell movement.

Notably, primary hCD4^+^ T cells exhibit pronounced compartmentalization during migration through dense 3D collagen matrices, with visible cortical ring structures indicative of active nuclear piston engagement **(Figure 4D)**. We posit that the nucleus dynamically interacts with multiple cortical rings at the bases of exploratory protrusions, and its eventual spatial repositioning governs the directional choice of migrating T cells. In structurally complex environments, T cells adopt complex exploratory configurations, such as trident-shaped or multi-compartment morphologies, to probe and explore multiple potential pathways. Conversely, during sustained unidirectional migration, they exhibit elongated, linearly organized bodies with stable compartmentalization, reflecting continuous hydraulic and contractile alignment **(Figure 4D)**.

Collectively, these results support a model in which increasing obstacle density amplifies the utility of the nuclear piston mechanism, allowing T cells to translocate their nucleus more effectively and enhance tissue infiltration and immune surveillance.

### Repolarization enhances migration of compartmentalized T cells

Next, we examined whether cell repolarization, which we model as a relocation of the leading edge with the expanding regions on each side, enhances T cells’ ability to navigate dense obstacle fields, particularly those forming impassable high-density discrete obstacle clusters. To test this, we simulated T cell migration within a randomized obstacle field **(Figure 5A-1, Movies 3-5)**, designed to resemble the configurations observed in 3D collagen matrix samples populated with human primary T cells **(Figure 5A-2)**. Here, we posit that the T cell’s ability to shift the direction of its leading segment expansion could facilitate a stirring effect, enabling the cell to escape from dead ends, *i*.*e*., promoting migration efficiency. To capture this behavior, we introduced a dynamic repolarization mechanism: after a fixed time interval Δ*t*_S*hift*_, the cell reorients its leading edge with the expanding regions on each side to probe a new direction. For this, we employed a tiered modeling approach, with the leading segment of the T cell exhibiting wide, narrow, and no repolarization capability **(See Methods)**.

**Figure 5.**
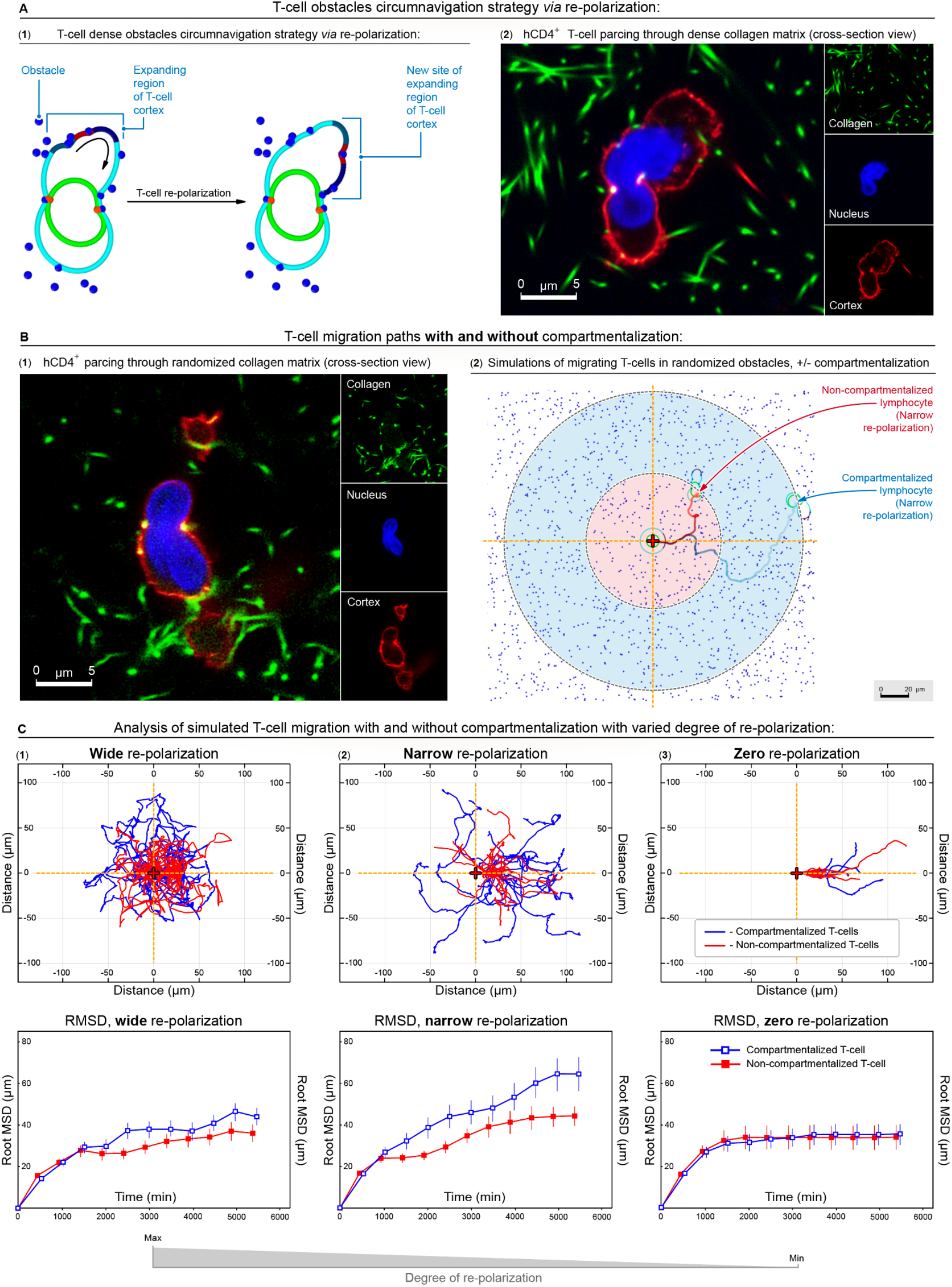
Computational analysis of the role of the T cell compartmentalization and repolarization as an explorative strategy to enhance the efficiency of the T cell migration in the randomized isotropic obstacle field. **A** - (**1**) Schematic visualization and (**2**) cross-section reference view of the primary hCD4^+^ T cell navigating within the complex obstacle field (collagen). Schematic (**1**) displays the explorative mechanism of the T cell migration *via* their leading compartment’s cortex repolarization. The T cell is shown as its 2D cross-section in a 3D collagen matrix for comparison with the 2D T cell model. **B** - (**1**) Cross-section view of the primary hCD4^+^ T cell passing through collagen of uneven density and (**2**) computational modeling of the T cell migration within the randomized unevenly dense obstacle field. T cell compartmentalization within the randomized obstacle field results in a more effective displacement, *i*.*e*., distance from the original position, compared to the non-compartmentalized T cell model. The simulations suggest that repolarization of the leading segment allows an optimized path-searching strategy to find a less confining, sterically permissive passage through the obstacle field. **C** - (*Top*) Individual T cell migration tracks within a randomized obstacle field: compartmentalized T cells (*blue*) cover a more significant displacement from their original positions than the non-compartmentalized T cells (*red*). (*Bottom*) Statistical analysis of migration efficiency, measured as a root of mean squared displacements (RMSD), in simulated T cells with and without compartmentalization under three ranges of leading segment re-polarization capacity, *L*^*Repol*^, defined as a measure of expanding region shift along the cortex: wide, narrow, and none. Wide re-polarization, *L*^*Repol*^ is equal to half of the cell circumference; narrow repolarization, *L*^*Repol*^ = *L*^*Front*^; no repolarization, *L*^*Repol*^ = 0. The results show that compartmentalizable T cells exhibit up to 90% greater migration efficiency than non-compartmentalizable T cells over the simulated time interval. Additionally, within both groups, re-polarization capability follows the same trend in enhancing migration efficiency, ranked from highest to lowest: narrow re-polarization > wide re-polarization > no re-polarization.

First, we focused on modeling T cells with an intermediate (narrow) repolarization shift capacity at < 74 degrees, to evaluate the specific contribution of the hydraulic nuclear piston mechanism (*i*.*e*., hydraulically sealed compartmentalization) to migration. To this end, we simulated individual T cell migration within a randomized obstacle field mimicking environments with alternating densely and sparsely packed 3D collagen **(Figure 5B-1)**. The resulting migration tracks **(Figure 5B-2, Movie 3)** demonstrate that over a fixed time interval (*t* = 6,000 min), compartmentalized T cells exhibit significantly greater displacement from their initial positions compared to non-compartmentalized counterparts. These findings are consistent with prior results **(Figure 1C)**, further supporting the idea that compared to the non-compartmentalized cells, compartmentalization facilitates more effective cell navigation through narrow interstitial gaps, such as those found in both simulated randomized obstacle fields **(Figures 5A-1 and 5B-2)** and collagen matrices **(Figures 5A-2 and 5B-1)**.

Second, with a large set of simulations, we explored how T cell displacement is affected by combining compartmentalization (hydraulic nuclear piston) with different tiers of leading compartment repolarization capacity: wide **(Figure 5C-1, Movie 4)**, narrow **(Figure 5C-2, Movie 3)**, and zero **(Figure 5C-3, Movie 5)**. Simulations of T cell populations (n = 20 per condition) reveal that narrow repolarization, when combined with compartmentalization, yields the highest root mean squared displacement (RMSD) from the original position **(Figure 5C-2)**. Wide repolarization with compartmentalization yielded the second-highest RMSD **(Figure 5C-1)**, whereas all other combinations, including non-compartmentalized T cells, exhibited lower RMSDs, which were not substantially different from each other **(Figure 5C-1 and 5C-3)**.

These results suggest that although wide repolarization might provide faster escape from dead ends, its cost in directional persistence leads to reduced cell displacement over long enough times. Therefore, optimal T cell migration in complex, obstructed environments requires a combination of hydraulic compartmentalization and a balanced repolarization capacity that is substantial enough to redirect around impassable regions, yet restrained enough to avoid inefficient, meandering random-walk migration.

### Uropod stabilizes the peristaltic polarity of T cells

As described above, in discretely crowded environments, such as reticular 3D collagen, the amoeboid T cells use peristaltic migration. One of the principal questions in this model is: how do the T cells maintain the stable polarity of peristaltic flow to ensure the directional persistence of migration? Here, we propose a biophysical model in which the uropod ensures the stability of peristaltic polarity. In this model, the uropod is the end-product of the peristaltic cycle, formed *via* the actomyosin contractility-driven collapse of the utilized part of the rear segment **(Figure 6A-1)**.

**Figure 6.**
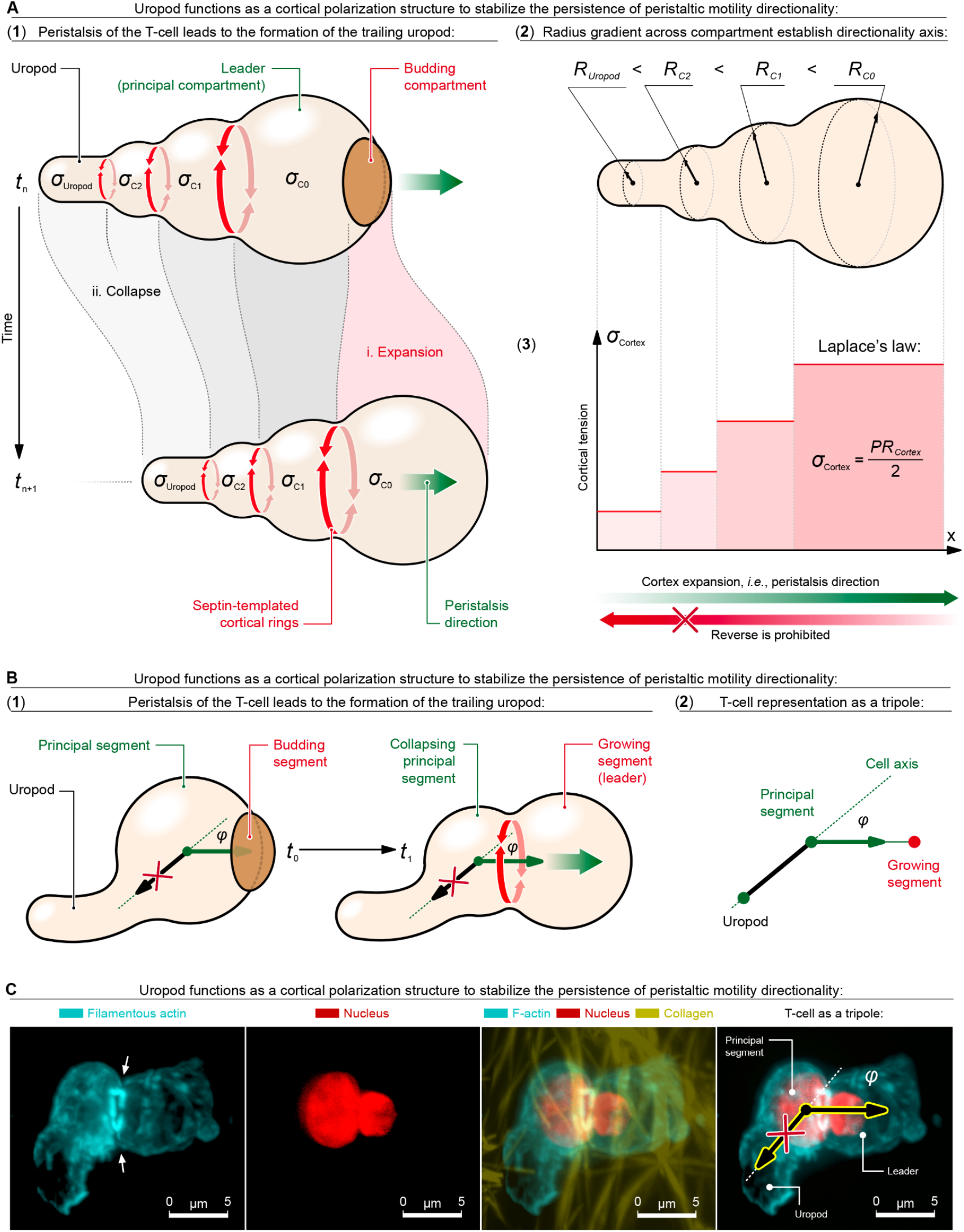
Uropod enhances the persistence of T cells’ amoeboid migration by establishing the polarity of peristalsis. **A** - (**1**) The T cell peristalsis-driven motility model represents a continuous migratory treadmilling: the T cell extends forward with a newly formed leader segment (expansion) while utilized segments contract, retract, and compact into the uropod (collapse), resulting in the T cell treadmill-like movement across the obstacle field. The cytoplasm and nucleus peristaltically translocate between the contractile segments towards the leader segment. (**2**) Variability of the sizes between the leading (expanding) and trailing (collapsing) T cell segments leads to the formation of the cross-section radii gradient. The T cell uropod features the smallest cross-section radius, resulting in a (**3**) least cortical tension σ_*Cortex*_ needed to sustain the cytoplasm hydrostatic pressure *P* per Laplace’s Law. **B** - The presence of the uropod ensures irreversible polarization of the peristaltic T cell, prohibiting a reversal of the peristalsis by re-expansion of the uropod into the leading (growing) segment. (**1**) The presence of a uropod renders the T cell into a 3-point system: uropod, principal segment, and budding/growing/leader segment. (**2**) Schema defining T cell as a 3-point system with the angular deviation φ between a newly formed growing segment and the current T cell axis (uropod-principal segment). **C** - 3D reconstruction of hCD4^+^ T cell within collagen matrix forming a minimal segmented configuration in agreement with the principal 3-point mechanical model, *i*.*e*., tripole: uropod, principal segment, and leader.

Since the uropod is the product of the structural compression of the pushing compartment, the cross-section radius *R*_*Uropod*_ of the uropod is generally smaller than the radius *R*_*c*_ of the other T cell compartments **(Figure 6A-2)**. Indeed, the uropod in various amoeboid lymphocytes is reported to feature a substantially smaller radius than the rest of the cell’s body ^47,82^.

According to Laplace’s law for hydrostatic systems, at any given moment, the expanding tension σ_*Uropod*_ within the surface layer of the uropod, induced by the cytoplasmic pressure *P* across all parts of the rear segment, is smaller than the cortical tension σ_*Cortex*_ of the remaining surfaces of the rear segment with larger radii **(Figure 6A-3)**. This structural arrangement ensures the irreversibility of uropod formation, preventing its re-expansion, as other segments experience greater cortical-expanding forces.

Laplace’s law provides the mechanical basis for the stabilization of peristaltic polarity in migrating T cells by the uropod. Specifically, the narrow uropod sustains the continuous collapse of the T cell’s trailing end, accompanied by the persistent formation of the large leader segments at the front end. Consequently, the compartmentalized T cells are biased towards acquiring the unidirectional treadmilling of cell segments and persistent amoeboid migration **(Figure 6A)**.

### Uropod increases the surveillance area during the random-walk migration of T cells

Stabilization of unidirectional peristaltic flow by uropod may decrease the ability of T cells to reverse the direction of migration and escape dead-end passages. However, the lack of backward motion could increase the persistence of the T cell migration and the area of immune surveillance. Moreover, deviations in the direction of forward movement could also affect the extent of the area surveyed by migrating T cells. To examine these possibilities, we model various degrees to which the uropod may constrain the T cell directionally and define the resulting surveillance area.

First, we considered the simplest representation of a T cell as a particle (centroid of the principal segment) that moves in the unconfined 2D space, *i*.*e*., in the migration field with no obstacles. We accounted for the uropod effect as the probability of angular deviation of the current migration steps with respect to the previous one. In this coarse-grained representation, the previous direction represents the orientation of the ‘uropod → cell body’ axis, and the new direction represents the ‘cell body → growing leader segment’ axis (**Figure 6B**). Notably, similar shapes are routinely observed for the amoeboid T cells (**Figure 6C**).

Second, we have set the limiting effects of the uropod at three levels, shown as the angular probability distribution curves for the directional deviations: (1) strict angular limitation **(Figure 7A-1)**, (2) intermediate angular limitation **(Figure 7A-2)**, and (3) low angular limitation **(Figure 7A-3)**. The results of these stochastic simulations using the T cell centroid **(Figure 7A, 1-3)** illustrate that limiting the angular distribution of the new direction relative to the uropod’s position significantly enhances the T cell’s ability to migrate further from the position of origin within the same number of the peristaltic cycles (*N*_*Cycles*_ = 1000) **(Figure 7B)**.

**Figure 7.**
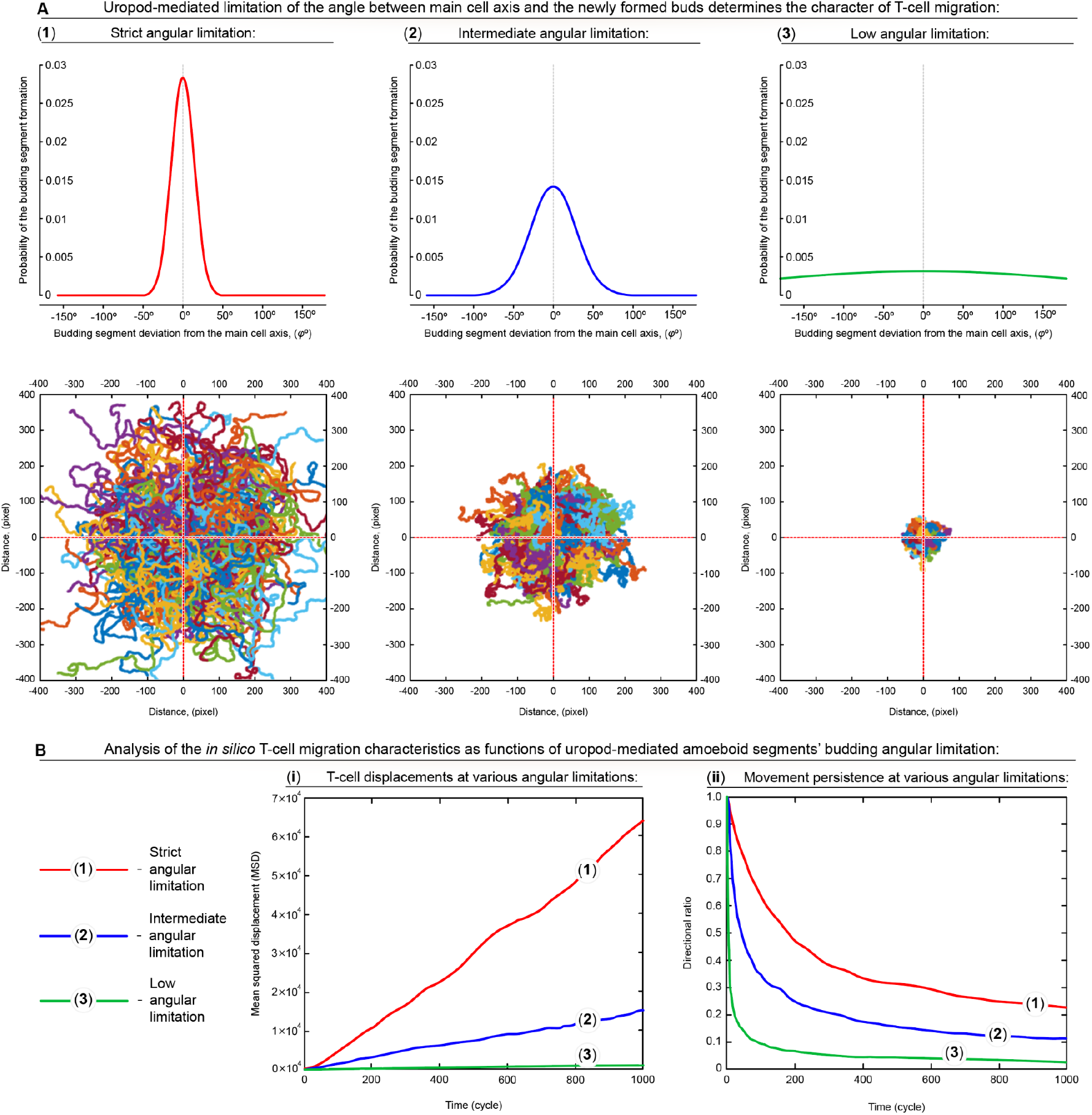
The degree of uropod-imposed constraints on the angular deviation of the T cell’s direction from its principal axis controls the efficiency of the lymphocyte’s random walk. **A** - **Top panels (1–3):** Distribution of angular deviations between the leading segment and the current cell axis for three conditions: (**1**) strict (narrow), (**2**) intermediate, and (**3**) low (wide) allowable angular deviation ranges. **Bottom panels (1–3):** Individual random-walk migration tracks of unrestricted amoeboid T cells (no obstacles) under the same conditions. A stricter angular deviation range (**1**) leads to a more efficient displacement from the point of origin, while a broader range (**3**) results in a reduced displacement. **B** - **Quantitative analysis:** (**i**) Mean squared displacement (MSD) and (**ii**) movement persistence of T cells across the three angular deviation conditions—strict (**1**), intermediate (**2**), and low (**3**). Higher restriction on angular deviations corresponds to increased movement efficiency and persistence.

### Uropod promotes confined T cell migration

Hereafter, we provide the details of modeling the interplay between the uropod-dependent restriction on the direction reversal and the geometric properties of the confining microenvironment. As a T cell model, here we use a three-segment system comprising the (1) uropod, (2) principal, nucleus-containing segment, and (3) newly formed, nucleus-receiving leader segment **(Figure 6B-1)**. The proposed structure can be simplified into a tripole, *i*.*e*., a three-point model **(Figure 6B-2)**. In a three-point model, the elementary cycle of the T cell amoeboid motility consists of three consecutive events: (1) generating the new leading segment (*i*.*e*., cell protrusion from the principal segment, (2) reassignment of the formed segment into a principal segment (*i*.*e*., the passage of nucleus through a constriction), (3) reassigning (*i*.*e*., collapse of) the former principal segment into a uropod. Additionally, to account for the uropod effects, we define the uropod as a structure that prohibits back steps by suppressing the probability of leading segment formation at the uropod’s location.

We use a discrete obstacle field to test the migration of the three-point T cell model. The discrete obstacle field closely approximates the reticular architecture of the lymphatic nodes ^10,41,93^, as well as the interstitial spaces ^10^, enriched in the dense reticular 3D collagen networks. In these environments, the individual collagen fibers partition the T cell’s cortex into the independently contracting interconnected segments, positioned within the obstacle-free spaces. Hereafter, we refer to these interconnected spaces as “chambers”. We postulate that T cells conform their morphology and peristalsis to the surrounding chambers, thereby minimizing the energy required for migration, specifically for the peristaltic flow of the nucleus and cytoplasm between adjacent chambers. To capture the structural complexity and the level of confinement of the reticular collagen networks in the simulation, we model the spatial positioning of the chambers as the elementary units (polygonal tiles) of free space between the ‘collagen obstacles’ where a migrating amoeboid T cell can fit its individual, septin-templated cortical rings ^20^.

Since the surrounding obstacles are discrete, they can form several openings around the T cells where the new leader segment can protrude **(Figure 8A-1)**. Subsequently, we subdivide the formation of the newly protruding segment into three steps: Step 1 - The protruding segment enters a neighboring chamber. The probability of choosing a chamber to enter is proportional to the size of the opening (gap) leading to this chamber. The direction of the uropod is excluded from choosing (no backsteps). Step 2 - The protruding segment either commits to the new chamber (which triggers translocation of the main body, *i*.*e*., principal segment, to this chamber, followed by updating the uropod direction) or retracts to the chamber with the main body. The coefficient *C* is a parameter of the model, such that if *C* = 0, the segment always commits to its first choice. If *C* > *C*_*max*_, the segment always retracts. If 0 < *C* < *C*_*max*_, both events are stochastically possible. Step 3 - If the protruding segment returns, it probes another neighboring chamber, repeating Step 2. Suppose all adjacent chambers are checked without commitment. In that case, the decision to move the main body to one of the checked chambers corresponds to the probability proportional to the largest openings in the ‘tested’ chambers. After step 3, the direction of the uropod gets updated. Note that the largest openings in the neighboring (new) chambers and the size of the openings in the current chamber are independent variables (see **Methods**). For these simulations, a T cell can be represented either as a three-point model with a uropod **(Figure 8A-2**, no backsteps) or as a two-point cell that can migrate from its current chamber in any direction **(Figure 8A-3)**.

**Figure 8.**
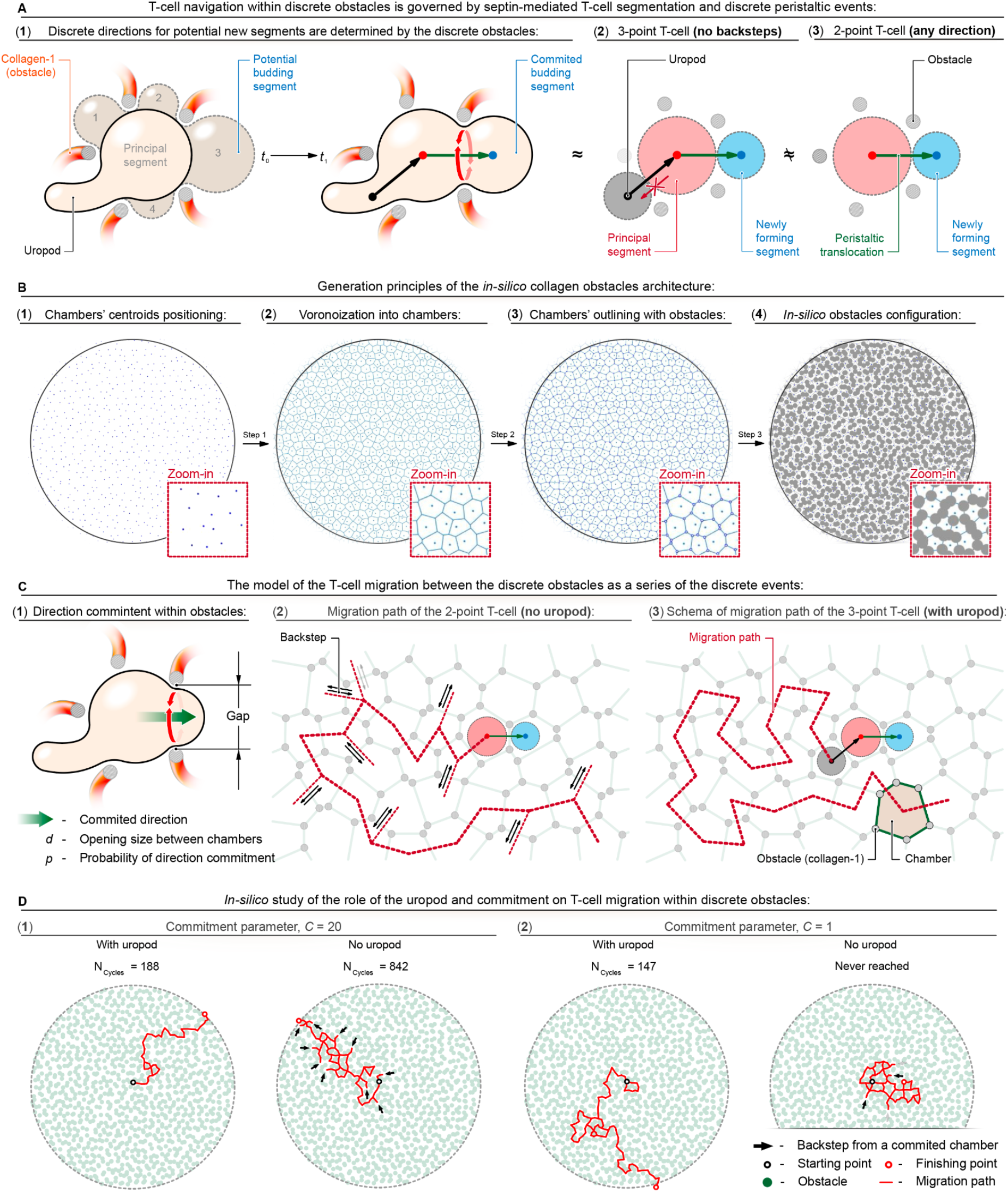
Schematic of the T cell migration modes in a discrete obstacle field (collagen fibers) with and without uropod. **A** - (**1**) The potential leader segment’s position and direction are determined by the geometric configuration of surrounding obstacles and the uropod’s position. (**2**) T cell with a uropod modeled as a 3-point system (no backstepping). (**3**) T cell without a uropod modeled as a 2-point system (allowing movement in any direction, including backstepping). **B** - Generation of the obstacle field mimicking a collagen meshwork: (**1**) Placement of chamber centroids; (**2**) Space partitioning into chambers using the Voronoi tessellation; (**3**) Positioning of obstacles at the vertices of the resulting polygonal chambers; (**4**) Defining obstacle radius. **C** - Schematic depiction of T cell migration and chamber commitment: (**1**) Schematic depiction of the T cell with uropod committing to the new chamber *via* the available passage, *d*, between the obstacles; (**2**) Migration of the 2-point T cell (without uropod) showing backstepping, as the leader segment can protrude in any direction, including revisiting previous chambers. (**3**) Migration of the 3-point T cell (with uropod) demonstrating unidirectional movement without backstepping. **D** - Simulated migration tracks of 2- and 3-point T cell models within the obstacle field (noise = 0.3; radius = 0.3, the whole field = 15 units): (**1**) Commitment parameter *C* = 20; and (**2**) Commitment parameter *C* = 1. *The probability of committing is proportional to the opening size d of the new chamber*. *The coefficient C is a model parameter: If C* = 0, *the segment always commits to its first choice. If C* > *C*_*max*_, *the segment always retracts. If* 0 < *C* < *C*_*max*_, *both events are stochastically possible*. Arrows highlight backstep events for the 2-point T cell model (without uropod). *N*_*Cycles*_ denotes the number of cycles used to reach the edge of the obstacle fields.

To generate the migration field with controlled geometric properties, we used an array of the chamber seeds **(Figure 8B-1)** to set the chamber configuration *via* the Voronoi tessellation algorithm **(Figure 8B-2)**. The seeds are placed based on the random perturbation of the hexagonal grid **(Supplemental Figure 1)**. This way, the extent of the perturbation (‘noise’ parameter) controls the distribution of the chamber sizes. The vertices of the resulting polygonal chambers were used to set the positions of the obstacles **(Figure 8B-3)**. This procedure was followed by setting the physical size (‘radius’ parameter *r*) of the obstacles that define the available size of the openings between the chambers (*i*.*e*., the level of confinement). The size of obstacles also defines the available space for the simulated T cells within the individual chambers **(Figure 8B-4)**.

Using the positional noise and radius as the two adjustable obstacle field parameters, we can generate a range of microenvironments with different geometries of the passage networks (labyrinths) for T cells to circumnavigate. Higher obstacle radius values narrow down the openings between the chambers, thereby mimicking a higher level of confinement and more significant deformation of the nucleus during migration. Higher noise values increase the complexity of the passage arrangement, characterized by a broader distribution of the number and sizes of openings between the chambers, including the formation of dead-end passages. For the quantitative analysis, we tested five levels of structural noise, defined as a fraction of the grid’s step: 0.1, 0.2, 0.3, 0.4, and 0.5, and three levels of the obstacle radius: 0.2, 0.3, and 0.4 **(Supplemental Figure 1)**.

We utilized the resulting obstacle fields with chambers to investigate the character of the T cell migration under various conditions. We compared the T cell models with the uropod (three-point model) and without the uropod (two-point model) to determine the effects of the uropod on T cell amoeboid navigation throughout the obstacle field **(Figure 8C)**. A three-point model of the T cell that enters the new chambers with the newly formed protruding leader segment is shown in **Figure 8C-1**. The examples of the migration path of the two-point (**Figure 8C-2**) and three-point (**Figure 8C-3**) T cell models demonstrate the backstep events as deviations from the main route for the two-point T cell model (**Figure 8C-2, ⇆**). The backsteps of the two-point T cell model decrease the efficiency of T cell migration, resulting in unproductive movement cycles. These unproductive movement cycles increase the number of steps required for the two-point T cell model to cover the same distance from the starting point as for the three-point T cell model.

In **Figure 8D**, we present simulation results comparing T cell migration with and without the uropod, highlighting how the chamber commitment parameter, *C*, modulates migration character in spatially heterogeneous environments. Simulations were conducted in an obstacle field defined by a spatial noise level of 0.3 units, an obstacle radius of 0.3 units, and a total field radius of 15 units (see **Methods**). Two- and three-point T cell models were analyzed under two distinct commitment regimes. In the low-commitment condition (*C* = 20, **Figure 8D-1**), cells exhibit extended exploratory behavior, sampling multiple neighboring chambers before committing to a migration path. Conversely, under the high-commitment condition (*C* = 1, **Figure 8D-2**), cells rapidly commit to one of the first accessible chambers, potentially bypassing a larger available ‘chamber’ or a more favorable (faster and more energy efficient) path. These simulations reveal how the presence of the uropod and the commitment parameter interact to shape the efficiency and directionality of cell migration through crowded, structured environments.

While we posit that T cells can select neighboring chambers with larger openings, such choices do not necessarily guarantee entry into a larger chamber. Indeed, information about a chamber’s size or the presence of additional exits becomes available only after that selected chamber is entered, *i*.*e*., explored by the protruding segment. As a result, small chambers with a single large opening (entrance) can act as traps for overly committed cells, particularly those that commit too early without sufficient environmental sampling. This justifies the exploratory migration strategy, which allows for flexible probing before making a directional commitment. In agreement with this notion, our simulations show that T cells equipped with a uropod under low-commitment conditions (*C* = 20, **Figure 8D-1**) successfully identify and follow the shortest path from the center to the periphery of the obstacle field. In contrast, under high-commitment conditions (*C* = 1, **Figure 8D-2**), T cells lacking a uropod often fail to reach the edge, highlighting the combined importance of directional flexibility and stable polarization in navigating complex environments.

Our data indicate that multi-chamber probing optimizes the T cell search of sterically permissive passages, and the uropod optimizes the spatial T cell surveillance.

### Uropod and exploratory commitment define the outcomes of spatial surveillance within tissue-like environments

The structure of the obstacle field creates a specific architecture of the passage network, constraining the directions of cell motion. This network architecture defines the character of T cell migration, similar to that of live T cell migration in the reticular collagen of varying density and structural anisotropy ^94,95^. **Figure 9A** shows a side-by-side comparison of the obstacle fields with the array noise = 0.5 and obstacles radius = 0.4 *vs*. the array noise = 0.3 and obstacles radius = 0.3.

**Figure 9.**
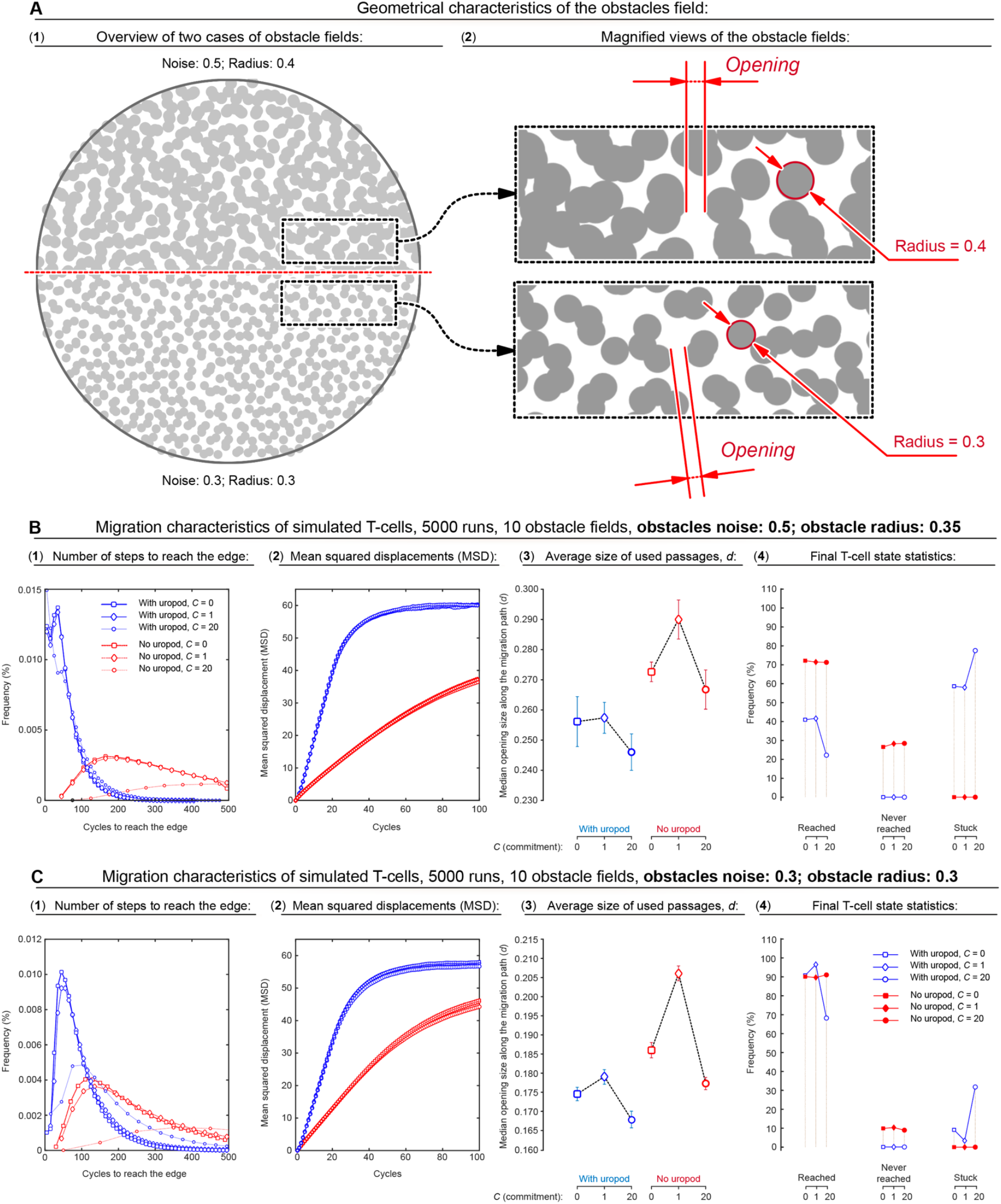
Quantitative analysis of the impact of the uropod, the obstacle characteristics, and the commitment parameter *C* on the simulated T cell migration. **A** - (**1**) Visual representations of the simulated obstacle fields: noise = 0.5, radius = 0.4 (*top*) and noise = 0.3, radius = 0.3 (*bottom*). (**2**) Magnified views of the obstacle fields, shown on panel **1**. **B** and **C** - Resulting, statistically analyzed migration characteristics of the simulated T cells from 5000 independent runs across 10 generated obstacle fields for each case. Results for the T cell migration in the obstacle fields with noise = 0.5 and obstacle radius = 0.35 are shown in **B**, while for the obstacle fields with noise = 0.3 and obstacle radius = 0.3 in **C**. (**1**) Simulation data for the number of cycles the T cells need to reach the obstacle field’s edge is shown as the frequency of cases for each number of cycles. The data are shown for T cells with and without the uropod, with the commitment parameters *C* set to 0, 1, and 20. *Note that the T cell with a uropod reaches the edge of the obstacle field with significantly fewer cycles than those without a uropod*. (**2**) Mean squared displacement of the T cells migrating with and without the uropod, with the commitment parameters *C* = 0, 1, and 20. *Note that the T cell with a uropod migrates farther from the original location than those without a uropod*. (**3**) Average size *d* of the utilized migration passages between the obstacles for the T cells with and without uropod, and with the commitment parameters *C* = 0, 1, and 20. *Note that the absence of a uropod allows backsteps, statistically allowing the T cell to utilize wider passages more frequently than those with the uropod. Note that at the commitment parameter C* = 0, *the T cells are prompted to always commit to the first chamber of choice. This results in the statistically narrower utilized migration passages on average for the T cells with C* = 0 *than those with C* = 1. *On the other hand, an extremely low commitment (C* = 20*) leads to frequent retractions and the resulting choices of new chambers with wider openings, even if that requires reaching them through narrow passages. This results in T cells displaying migration with a statistically higher percentage of transitions through narrow openings, while C* = 1 *case better balances the size of available openings in the current and the new chambers*. (**4**) The final states of the T cells migrating throughout 5000 cycles are split into three groups: T cells that reached the obstacle field’s edge, T cells that never reached the edge, and stuck T cells. *Note that the T cells with the uropod do not fall into the ‘never reached’ group. Instead, they either reach the edge or get stuck due to the uropod enhancing their migration persistence while disabling their ability to backstep from the trapping (dead-end) chambers. Note that T cells without the uropod never get stuck due to their ability to backstep from the trapping chambers. Also, note that T cells with the uropod get stuck more frequently in the obstacles with higher noise due to the higher number of dead-end chambers generated by the deviation of the obstacle positions from the ideal hexagonal grid*.

To quantitatively assess the role of the uropod and the commitment parameter *C*, we employed a large-scale statistical simulation approach. For each experimental condition, we generated 100 independently randomized obstacle fields and conducted 5,000 T cell migration simulations per field, resulting in a total of 500,000 simulations per condition. We examined three levels of commitment behavior (*C* = 0, 1, 20) across two structurally distinct environments: (1) high-noise fields with obstacle radius 0.35 and spatial noise of 0.5 **(Figure 9B)**, and (2) low-noise fields with obstacle radius 0.3 and spatial noise of 0.3 **(Figure 9C)**. In all simulations, the obstacle fields were circular with a fixed outer radius of 15 units. This high-throughput approach enabled the extraction of robust average behaviors and migration outcomes across diverse microenvironmental configurations.

We focus on four key output parameters (**i-iv**) of the T cell amoeboid migration: (**i**) the number of steps to reach the edge of the obstacle field **(Figures 9B-1** and **9C-1)**, (**ii**) mean squared displacement, MSD, from the original location **(Figures 9B-2** and **9C-2)**, (**iii**) averaged size of the used openings along the entire migration path of the T cell, *i*.*e*., until T cell reaches the edge of the obstacle field or the end of the simulation run (*N*_*Cycles*_ = 500) **(Figures 9B-3** and **9C-3)**, (**iv**) the percentage of the simulated T cells that reached the edge, never reached the edge by the time of the run, and stuck within the impassable obstacle configurations **(Figures 9B-4** and **9C-4)**.

Simulation results show that T cells with uropod reach the edge of the obstacle field within significantly fewer cycles than those without the uropod, as observed for both types of tested obstacle fields **(Figures 9B-1** and **9C-1)**. These data agree with the substantially higher migration efficiency of the simulated T cells with the uropod, as indicated by the MSD **(Figures 9B-2** and **9C-2)**. Notably, simulated T cells without the uropod experience a significantly greater slowdown in reaching the edge of the obstacle field as the commitment parameter (*C*) increases from 0 or 1 to 20 (see cycles to reach the edge in **Figure 9B-1** and **9C-1**). This effect is also observed for T cells with the uropod in obstacle fields with low noise (0.3) and smaller obstacle radius (0.3) **(Figure 9C-1)** compared to those with higher noise (0.5) and larger obstacle radius (0.35) **(Figure 9B-1)**. The underlying reason for this loss of passage efficiency is the greater availability of wider openings between chambers in low-noise/small-radii obstacle fields, which increases the time that cells can explore local areas. As a result, T cells with the low-commitment parameter (*C* = 20) take longer to navigate through low-noise, small-obstacle-radius fields than through high-noise, larger-obstacle-radius environments with fewer options to explore. For T cells without a uropod, a low-commitment strategy is always slow because they can reverse direction, and even with fewer options to explore in high-noise/large-radii fields, they lose time on futile back-and-forth runs.

**Figures 9B-3 and 9C-3** illustrate the analysis of the average opening size used by T cells along their migration path for high-noise, larger-obstacle-radius environments **(Figure 9B-3)** and low-noise, small-obstacle-radius environments **(Figure 9C-3)**. These data provide a comparison of T cell migration with and without a uropod, as well as across different commitment parameters (*C* = 0, 1, and 20). Notably, T cells lacking a uropod exhibit backsteps, which statistically increases their likelihood of utilizing larger passages more frequently than T cells with a uropod. When the commitment parameter is set to *C* = 0, T cells are forced to commit immediately to their first-choice chamber, resulting in a statistically narrower migration passage on average compared to T cells with a lower commitment, *C* = 1. Alternatively, an extremely low commitment level (*i*.*e*., *C* = 20) results in stalled commitment decision-making, frequent leading segment retractions, and prolonged residence in chambers with many openings, ultimately choosing a new chamber with another large opening but not necessarily a large opening to reach this chamber from the current position. As a result, T cells with *C* = 20 display sub-optimal migration along trajectories with both broad and narrow passages.

Finally, we categorized the outcomes of simulated T cell migration in both high-noise, large-obstacle-radius environments **(Figure 9B-4)** and low-noise, small-obstacle-radius fields **(Figure 9C-4)** into three distinct groups: (**1**) T cells that successfully reached the edge of the obstacle field, (**2**) T cells that remained active but never reached the edge within the simulation time frame, and (**3**) T cells that became mechanically stuck. Notably, T cells lacking a uropod were observed in the ‘never reached’ group but never became irreversibly trapped, due to their fully preserved ability to backstep from the trapping chambers. In contrast, T cells with the uropod were never found in the ‘never reached’ category. Instead, they either reach the field’s edge or get stuck due to the uropod, assuming it completely suppresses T cells’ ability to backstep from the dead-end (single opening) chambers. Moreover, T cells with a uropod become stuck more frequently in fields with higher noise, due to the higher number of dead-end chambers. The dead-end chambers are generated by the significant displacements of obstacles from the regular grid, resulting in the overlap of neighboring obstacles that effectively close all openings except the one that cells can use to enter. These structural dead ends reduce the migration paths and elevate the risk of irreversible trapping when the uropod impairs reverse migration.

Together, our results indicate that the presence of the uropod could optimize the T cell migration and its ability to choose a less sterically restrictive path, leading to energy conservation, faster migration, and more efficient immune surveillance.

## DISCUSSION

The lack of mechanistically predictive models of amoeboid migration in tissues ^29,45^ impedes our ability to counteract cancer metastasis and limited infiltration of immune cells, such as cytotoxic NK cells and CD8^+^ T cells, into solid tumors. To facilitate the understanding of amoeboid migration, which is critical for effective immune surveillance ^96–98^ and cancer immunotherapies ^99–103^, we explored the biophysical cooperation between the septins, actomyosin cytoskeleton, nucleus, and the surrounding obstacles. Our data suggest that these components act synergistically to form a hydraulically driven nuclear piston at the sites of collisions between the cell cortex and the extracellular matrix **(Figures 1B-1, 2A**, and **2B)**.

Specifically, with the assistance of F-actin-septin curvature-sensitive interactions and cytoskeletal condensation ^20^, the T cell recognizes and responds to the indenting cortical deformations by forming tight hydraulic seals around the passing nucleus. The resulting seals create a condition for developing a difference in hydrostatic pressure between the cytoplasmic segments on each side of the nucleus ^20,32,59,64^, which becomes a part of the hydraulic piston. The formed hydraulic piston allows the rear segment of the T cell to accumulate energy in the form of hydrostatic pressure and cortical tension. The accumulated energy powers forward thrust on the nucleus, resulting in sustained movement within the confining environment.

Moreover, our proposed septin-templated hydraulic piston model suggests that a high density of obstacles strengthens the nuclear piston mechanism, allowing T cells to push their nucleus more effectively, thereby improving infiltration and immune surveillance. Mechanistically, the nuclear thrust is based on the pressure difference between the front and rear compartments necessary for forward propulsion of the nucleus. According to the suggested conceptual framework that motivated our computational models, this pressure difference may arise from the presence of the uropod with high actomyosin contractility and small radius ensuring the persistence of a high pressure at the rear compartment during the motion of the nucleus and the changes in cell shape **(Figure 2C)**.

Therefore, our results suggest that the uropod is not simply a byproduct of the rear cortex collapse upon its contraction or a reservoir for molecules involved in signal transduction, cell-ECM, or cell-cell interactions but an important component of the cell engine: the physical mechanism enabling efficient and stable (unidirectional) force generation to propel the nucleus through constrictions.

Intriguingly, this persistent nuclear thrust is enabled by the spatial positioning of the cortical hydraulic seal, which forms at obstacle-induced indentations on the T cell surface (as shown in **Figure 1B-1** and previously reported ^20^). This implies that the thrust mechanism operates precisely where it is most needed, at sites of mechanical impedance where the nucleus faces resistance. Based on our presented data and simulation results, we propose that septin-templated seals represent an evolved feature of amoeboid motility, enabling the generation of intensified, hydraulically transmitted forces to drive efficient nuclear translocation. Importantly, this hydraulic effect appears to occur precisely where it is most needed - at sites of local geometric constraint imposed by the extracellular matrix. Future investigations should explore whether this hydraulic “booster” is regulated or modulated by cytoskeletal dynamics, such as actomyosin contractility or septin-mediated reinforcement at the seal, and whether it plays a broader role in confined migration across other immune or cancer cell types.

Furthermore, despite the reported migration-interfering effects of the nucleus due to its large size and high rigidity ^44,48^, the septin-enabled hydraulic effects may effectively transform it from a bulky burden into an essential part of the mechanism driving amoeboid locomotion of T cells under confinement.

Historically, the first conceptual elements of the hydraulically active nuclear piston have been reported in non-immune cells. Namely, the linker protein nesprin can provide a direct mechanical bond between the nucleus and the actomyosin cortex in primary fibroblasts. This mechanical connection enables actomyosin-driven forward nucleus traction, generating frontal cytoplasm pressure that hydrostatically expands the cell’s front into lobopodial protrusions ^64^. In the mesenchymal stem cells, this mechanism is amplified by pressure-sensitive ion channels. These channels, upon sensing nuclear piston-induced pressure, enhance osmotically driven expansion of the lobopodial protrusions ^50^. Collectively, these studies describe linker-based nuclear piston systems. The linker-based system utilizes mechanical traction to drive nuclear propulsion, facilitated by molecular adapters that transmit actomyosin-generated force directly onto the nucleus, thereby generating hydrostatic pressure.

Additionally, the linker-based nuclear pistons require external adhesions to the extracellular matrix, which serve as mechanical anchors or fulcrums, enabling traction for nucleus translocation. As a result, for a mesenchymal cell, the migration process is typically accompanied by considerable frictional resistance, created between the moving nucleus, cortex, and the anchoring environment. In contrast, the adhesion-free amoeboid T cells seem to employ a fundamentally different motility mechanism: a purely hydraulic nuclear piston that relies solely on sliding by pressure difference between cellular compartments to drive nucleus movement. This amoeboid mode of nuclear propulsion does not require external adhesion or molecular linkers for mechanical pulling. Instead, it operates through steric interactions between the T cell and its surrounding environment **(Figures 1A** and **1B-1)**, mediated by septin-templated cortical rings ^20^. The steric cues, which are sufficient to induce pressure compartmentalization, support a sustained piston-like thrust of the nucleus through the confining spaces.

Notably, the obstacle-induced cytoskeletal changes that assist T cell nucleus propulsion have also been described for dendritic cells, where the local narrowing of environmental spacing triggers cell cortex adaptation in the form of Arp2/3-mediated F-actin polymerization and cortex densification, which are suggested to soften the nucleus and facilitate its passage ^59^. However, whether this mechanism renders the dendritic cell nucleus into a hydrostatically compartmentalizing piston remains unknown. The role of alternative force-generating mechanisms, such as microtubules (MTs) and MT-associated motor proteins, in the confined cell migration described above is also unknown. Specifically, it has been suggested that dynein and kinesins are crucial for cell invasion ^104^ and long-distance force transmission across the elongated cell body *via* microtubules, which act as non-stretchable cables ^104,105^, preventing the rupture of overstretched cells. These properties may be necessary for dendritic cells, which are mechanobiologically and phenotypically distinct from T cells ^106^. For example, in tight confinement, *i*.*e*., during diapedesis, highly deformable dendritic cells maintain their structural integrity under extreme deformations, such as overstretching, thinning, branching, or a tug-of-war state ^32^. Therefore, it remains to be determined whether T cells and dendritic cells habitually share the hydraulic nuclear piston or other mechanisms that allow dendritic cells to generate and coordinate a balance of pushing-pulling forces to invade tight confining spaces^18^.

The discussed concept of the hydraulically driven nuclear piston inherently complements the mechanosensory role of the nucleus suggested for cells migrating through confining environments. Specifically, nucleus deformations induced by collisions with surrounding obstacles can cause calcium ions to be released from the nucleus, thereby activating actomyosin cortex contractility, which helps the nucleus overcome the confining obstacle ^54^. Since the degree of Ca^2+^ release is proportional to the degree of nucleus deformation, which is reversely proportional to the nucleus’s mechanical rigidity, the nucleus size and mechanical properties would be among the key characteristics that define the migratory properties of the cell. Thus, lymphocytes can tightly integrate the mechanosensory and hydraulic roles of the nucleus into a decision-making unit that operates during migration within confining environments. In the future, this conjecture can be tested both *in silico* and *in vivo* by targeting calcium ion flux or by altering nuclear deformability.

The second major part of our work is the characterization of the uropod contribution to the efficiency of T cells’ circumnavigation in complex, confining environments. Using a stochastic agent-based modeling approach, we demonstrated that the uropod of migrating T cells is not only a part of the nuclear thrust mechanism as described above but also a key contributor to the T cells’ migration strategy, relying on the stability of the amoeboid cell polarization and migration persistence. Thus, we argue that the uropod enhances T cell migratory efficiency during spatial surveillance within tissue-like environments. As with nuclear propulsion, the physical basis behind the stabilization of T cell polarization by the uropod is its relatively high cortical tension and small size, preventing expansion under high hydrostatic pressures at the rear of the cell (in accordance with Laplace’s law) **(Figure 6A-3)**. This structural adaptation ensures the irreversibility of the uropod formation, stabilizing polarization of T cells.

Our results also suggest that control over the stability of the T cell’s uropod is a potent strategy for re-engineering T cell motility to tune its parameters, such as directionality and persistence, and modulate the T cell’s ability to reverse its migration direction to avoid dead-end traps in dense solid tumors. Specifically, our results suggest that T cells without the uropod are never trapped in simulated tissue-like environments. In contrast, while T cells with the uropod can become trapped in dense tissues, they exhibit enhanced spatial surveillance. Indeed, our results suggest that the stability of the uropod may influence T cell migration and its ability to select a less sterically restrictive path, thereby leading to energy conservation, faster migration, and more efficient immune surveillance. Thus, effective T cell migration requires both compartmentalization and a balanced repolarization ability to stir and navigate around an impassable density of obstacles, avoiding the ineffective random-walk modality of migration. In the future, this conjecture could also be tested experimentally by targeting the uropod stability, *e*.*g*., *via* activation of the TCR, suppression of the uropod cytoskeleton’s crosslinkers, regulation of the septin activity, and control over the actomyosin contractility.

In summary, this study establishes the feasibility of the nuclear piston mechanism as a means to enhance immune cell migration under tightly confining extracellular conditions. Although based on coarse-grained representations of cell behavior, our findings support the potential for a **multiparametric control strategy** in which modest, incremental changes across several key parameters can collectively produce substantial effects on migratory efficiency. These parameters may include actomyosin contractility, septin-mediated formation of hydraulic seals, and nuclear rigidity. Adopting a mechanistic framework for modeling immune cell dynamics opens the door to computer-aided engineering of T cell locomotion in complex tissue environments. Ultimately, the development of a broadly applicable computational platform for mechanistic analysis and target identification could enable the **rational reprogramming of immune cell migration**, allowing for optimized infiltration, sustained activity, and improved immune surveillance in the hostile microenvironments of solid tumors and aging tissues.

## METHODS

### Bead-spring cell model accounting for the biomechanics of cortex and nucleus

A 2D bead-spring model was implemented to evolve the cell and nucleus boundaries according to passive forces that depend on cell shape, as well as motor-driven active forces (**Figure 1**). These boundaries are represented as two closed linear chains of beads connected by springs ^107,108^. The parameters utilized in this simulation are tabulated in **Supplemental Table 1**. Forces consist of stretching, bending, area conservation, excluded volume, nuclear-centering, and contractile forces. The sum of all forces on each bead, *i*, that make up the cortex and nucleus, is used to evolve the positions of each bead r _*i*_, over time, according to the following equations ^107^:

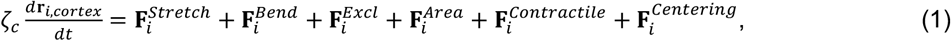

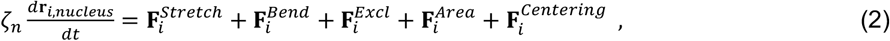

where ζ_*c*_ and ζ_*n*_ correspond to the effective drag coefficient for the cortex and nucleus, respectively.

The stretching and bending forces on the cell cortex beads represent the elastic response of the combined plasma membrane and adjacent actin cortex to stretching and bending, and similarly for the nucleus. Assuming cytoskeletal mechanics dominates both, we use the same reference values for both cell cortex and nucleus. The stretching force is:

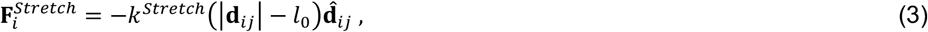

where *k*^*Stretch*^ is the spring constant (which depends on time; see below), *l*_0_ is the equilibrium length of the bead separation *d*_*ij*_ = |r_*j*_ − r_*i*_|, and 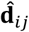 is the unit vector from bead *i* to bead *j*. The bending force is:

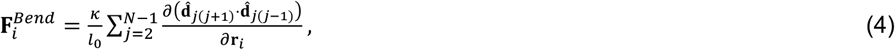

where *k* is the bending rigidity for the cell cortex or nucleus.

Excluded volume forces act between the cell membrane and the nucleus when bead pairs of segments α and β overlap. The force on α is exerted along the direction of vector d_α β_, which joins the closest approaching points between the two pairs ^107,109^:

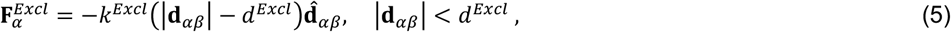

and, similarly, for β. This force is distributed to the beads at the end of the bond segments in proportion to their distance from the closest point. This force ensures that the nucleus stays inside the cell (*i*.*e*., the boundaries of the cell and the nucleus do not cross each other). The excluded volume force is the main force contribution that limits the timestep in simulations to reduce such unphysical bond crossings.

### Area conservation

We impose an energy cost for the deviation of the cytoplasm-containing area from a reference area *A*_0_ to conserve the area between the cell and nucleus boundaries. The energy and corresponding forces on the beads on the area boundaries are:

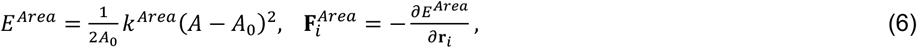

where *k*^*Area*^ is the effective area spring constant. The reference area *A*_0_ is the area of the region between the cell cortex and nucleus at the start of the simulation. The same expression was used as an energy cost for the area within the nucleus, using the same *k*^*Area*^ and a reference area equal to the nuclear area at the start of the simulation.

The area force is not applied to nuclear or cortex beads at a distance less than *d*^*Excl*^ away from each other, representing the absence of cytoplasm between the nucleus and cortex, at locations where they approach.

### Blebbing and actomyosin contractile force

We assume that the simulated cell is polarized with a contractile back region and a front region of length *L*^*Front*^, which consists of the leading edge and two flanking expanding regions that form blebs. Specifically, the front region is divided into the leading edge of length 0.28 *L*^*Front*^ and two flanking regions of length 0.36 *L*^*Front*^ each (see **Figure 1D**). The flanking regions are assumed to alternate in blebbing, representing the fluctuating bleb regions of motile T cells. Since blebs are plasma membrane regions disconnected from the actomyosin cortex, we represent them as a weakening of the simulated cell boundary stretching stiffness in Eq. (3). Bleb expansion and retraction is simulated as oscillations of stretching stiffness:

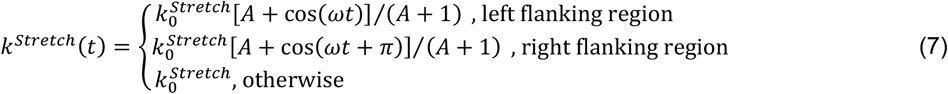

where *A* > 1 is a coefficient that determines the minimum and maximum values of *k*^*Stretch*^(t) and 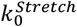is the reference spring constant.

Actomyosin contraction is represented as an inward force in the direction of unit vector 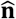 normal to the cell boundary applied to all cell cortex beads, excluding beads in the leading-edge segment:

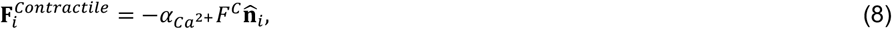

where *F*^C^ defines the basal force magnitude. Recent studies suggest that nuclear deformations may trigger increased cortical tension through the release of calcium ions (Ca^2+^) ^54,55,110^. To account for this effect, we use the nuclear area force as a proxy of nuclear deformation by including a parameter α_*Ca* ^2+^_, which has a value larger than unity (α_*Ca* ^2+^_ = 1.5) when the nuclear area force exceeds a threshold of 3*F*^C^ and is equal to 1 otherwise.

We further assume a reduction of contractile force to zero when the cortical boundary closely approaches the nuclear boundary (cortical beads reach a distance less than *d*^*Excl*^ from nuclear beads), consistent with decreased phospho-myosin intensity in T cell cortical regions near the nucleus ^20^.

### Nuclear centering

To represent the centering of the nucleus to the central region of the cell by organelles and microtubules ^83,84^, a weak spring centering force was included along the direction of the center of mass of the nucleus beads, r^*C*OM,n^, and cell cortex beads, r^*C*OM,n^:

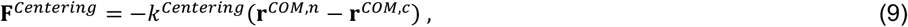

where *k*^*Centerin*g^ is a spring constant. This centering force is distributed equally to each bead of the nucleus by dividing the force by the number of nuclear beads. An equal and opposite force is distributed equally to each cell cortex bead.

### Extracellular matrix obstacles

In our 2D model, the extracellular matrix (ECM) is represented by obstacle beads fixed in space. Excluded volume forces between obstacles and the cell membrane are calculated similarly to Equation 5.

To study how cell migration depends on ECM density, we created a grid of obstacles with increasing density along the x-axis (**Figure 4A-1**). The grid is a lattice with vertical and horizontal distances between obstacles decreasing according to *dy*_n_ = *f*_n_(*dy*_n_ − *d*^*Excl*^) + *d*^*Excl*^ and 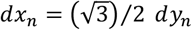, where *f*_n_ = 1 − *n*ε. Here *n* represents the column index, ε is a small parameter compared to unity (ε = 0.025), and *dy*_0_ was set equal to the diameter of the initially circular nucleus. The vertical shift of each row was chosen from a uniform probability distribution between 0 and *dy*_n_ for the first obstacle in each column. Subsequent obstacles in the corresponding column were placed at regular intervals.

In another set of simulations, we studied motion in an inhomogeneous environment by placing obstacles at random locations according to a uniform probability distribution around a simulated cell placed in the middle of a simulation domain.

### Compartmentalization by septin-templated cortical ring

A primary assumption in this model is that cytoplasmic fluid flow around the nucleus does not limit cell motility. However, forming a septin-templated ring results in a seal, which abruptly limits cytoplasmic flow. We expect the flow rate of cytoplasmic fluid through the narrow passageway between the cell membrane and the nucleus to depend strongly on passage width.

Implicit compartment boundaries were added to simulate septin-templated ring formation at regions where the nucleus is very close to the cell boundary due to nearby ECM obstacles. We identified candidate compartment boundary cortex (CBC) and compartment boundary nucleus (CBN) beads by checking every time step and finding instances where a cell cortex bead is close to both an obstacle and a nuclear bead (below a threshold distance of 1.3 μm and selecting the cortex and nuclear beads with the smallest distance). An implicit seal was formed when two CBCs formed at a distance > 2.6 μm from each other along the contour of the cell cortex. To account for the fact that the septin ring has a finite lifetime and is stationary with respect to the collagen fibers, once an implicit seal was formed, the identities of the CBC and CBN beads were unchanged for a lag time τ_*lag*_, after which we re-check for sealing. The CBC and CBN beads were allowed to move with the same forces as other beads during τ_*lag*_. The numerical value of τ_*lag*_ was tuned such that CBC and CBN neither slide away far from the constriction (too large τ_*lag*_) nor do they exhibit multiple back and forth jumps between neighboring beads (too short τ_*lag*_) during nuclear passage between obstacles.

When a seal forms, the system is divided into three conserved areas: behind the nucleus (C1), inside the nucleus (N), and in front of the nucleus (C2) (**Figure 1C-2**). The reference areas for C1 and C2 in Equation (6) were equal to the initial reference area *A*_0_ prior to compartmentalization, multiplied by the fraction of the whole area between the cortex and cell nucleus at the instant of compartmentalization. The reference area for the nucleus remained the same. When the seal is broken (after the cell passes through narrow confinements) the C1 and C2 compartments are merged.

### Repolarization

T cells must move through complex environments that would require them to change directions during amoeboid migration. It has been observed that each T cell locomotion cycle includes movement of the nucleus and cytoplasm into the front region of the cell while the back region assumes the uropod shape which is suggested to prevent repolarization of amoeboid T cells ^20^. To simulate cell turning and allow for small changes in direction, we subjected the leading-edge bead assignment to a shift across the rest of the cell cortex beads by *L*^*Repol*^ in either direction during a fixed time interval of the simulation. The leading-edge assignment was allowed to shift to either the left or the right of its previous position on the cortex. Since T cells move to avoid highly confining environments ^20,67^ the shift in bead assignment would endure for a fixed time interval, Δt_S*hi*ft_, after which the leading edge assignment shifts to other cortex beads in either direction. This leading-edge reassignment to different beads of the cell cortex causes the cell to change the direction of migration. Applying this mechanism leads to simulated T cell turning and changing direction by small amounts. We examined three cases: wide repolarization where *L*^*Repol*^ is equal to half of the cell circumference; narrow repolarization, *L*^*Repol*^ = *L*^*Front*^; no repolarization, *L*^*Repol*^ = 0.

### Parametrization of the obstacle field for modeling uropod-dependent migration strategies

To generate an obstacle field for the coarse-grained model T cell migration, we start with a regular hexagonal grid with 30 rows and 30 columns of points. The columns are separated by 1 unit, while rows are separated by √3/2 units. Each other row shifts by 0.5 units to achieve the hexagonal arrangement. Then, the positions of each point are randomly perturbed in x and y directions by a uniformly distributed noise with a magnitude serving as a parameter of the obstacle field generation, ‘noise.’ Next, each point is used as an input (seed) for the Voronoi tessellation and Delaunay triangulation. The built-in MATLAB function ‘voronoi’ returns the coordinates of the Voronoi vertices and the indices of the Voronoi cells. The function ‘delaunay’ returns a three-column matrix where each row contains the index of the input points that make up a triangle. Using this information, we find indices of the neighboring seeds (chambers) and the length of the Voronoi cell edges *S*_*ij*_ between each pair of seeds, *i* and *j*. The obstacles are assumed to be positioned at the Voronoi vertices, while the T cell segments move discretely from one chamber seed to another neighboring one through the corresponding Voronoi edge. After we define the obstacle size as another parameter of the field generation ‘radius,’ (*r*) we treat the space of that radius around each vertex as an inaccessible area for the T cell. Therefore, the value *S*_ij_ serves as the size of the opening between the chambers *i* and *j*. If *S*_*ij*_ < 2*r*, this opening is considered blocked so that the T cell cannot pass between chambers *i* and *j* through this Voronoi edge (*i*.*e*., chamber wall). A T cell is considered to reach the edge of the obstacle field when the position of its principal segment (with the nucleus) becomes over 15 units away from the center of the field where T cells start their migration runs.

### Uropod-dependent migration model

Our coarse-grained three-point T cell model represents the migration process with a peristaltic cycle that includes exploratory protrusion into available spaces (that we refer to as chambers), pushing the nucleus through the septin-templated cortical rings at the collagen indentations (that we refer to as openings between the chambers), and retracting the uropod to the space previously occupied by the cell body. In the model, the T cell can invade into any neighboring chamber through an opening, if its size *S* > 2*r*. However, the probability of choosing a specific opening is proportional to its size. For example, if the principal segment (second point of the three-point cell model) is in the chamber with *k* such openings of size *S*_k_, first a random opening, *i*, is selected by the MATLAB ‘randi’ function. Then, if *S*_i_/∑_k_ *S*_*k*_ < *R*, where *R* is a random number from a uniform distribution between 0 and1, the protruding segment (the first point of the three-point cell model) moves to this chamber. Otherwise, another random index, *i*, and another random number, *R*, is generated. The process is repeated until the condition is satisfied and the chamber is chosen for protruding. The model with the uropod excludes from choosing the chamber where it is located so that the protruding segment cannot move to the chamber with the uropod (which would represent a back step because the uropod occupies the principal segment location from the previous cycle). If no openings are available except for the back step (*k* = 1, dead-end chamber), the T cell with the uropod is considered stuck, and the simulation is terminated.

Otherwise, after the movement of the protruding segment into a new chamber, the algorithm checks if 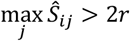 and 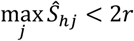, where *Ŝ*_*ij*_ is the size of the opening *j* in the new chamber *i, C* is the commitment parameter, and *R* is a random number between 0 and 1. The first condition checks if the new chamber has any openings other than the one just used, and the second condition stochastically accepts or rejects the new chamber for nucleus propulsion (*i*.*e*., the cell commits to complete transmigration between the chambers). This way, the probability of accepting the new chamber with available openings depends on the value of *C*. If *C* = 0, the second condition is always satisfied, and the chamber is always accepted. With increasing *C*, the probability of rejecting the chamber also increases because the randomly generated value of *R* needs to come out smaller than 1/C. If the chamber is rejected, the protruding segment retracts (the first point returns to the location of the second point), and the algorithm repeats the selection and commitment trial among the remaining *k* − 1 options. When no new chamber is accepted (expected for large values of parameter *C*), the commitment is ‘enforced’ based on the values of the largest opening in the new chambers 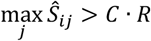. This selection is done in the same way by randomly selecting a chamber *h* and checking the condition *max* 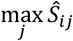, so that new chambers with larger openings are prioritized. In cases when all 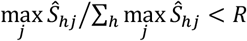, *i*.*e*., all neighboring chambers are dead-end chambers, the T cell with the uropod is considered to be stuck, and the simulation is terminated. The cell without the uropod can always go back and find an alternative path, or keep moving in circles, or back-and-forth. In other words, without the uropod, a T cell can be locally trapped but cannot be stuck the same way as a cell with the uropod that can stop moving.

Once the protruding segment commits to a new chamber, the positions of the three-point model are updated so that the principal segment (second point) moves to this new chamber and ‘joins’ the first point, while the uropod (third point) moves to the vacated old chamber. This completes the cycle, and the process of protruding, retracting, committing, and moving is repeated so that the T cell continues to move from chamber to chamber until it becomes stuck, reaches the edge of the obstacle field, or exhausts the simulation time (500 cycles).

The commitment strategy in our model is defined based on the control parameter *C* and the sizes of the inter-chamber openings rather than on the neighboring chamber sizes. Our rationale is that, in this study, we consider the formation of the septin-templated cortical rings, which provide a hydraulic seal to boost nuclear passage through narrow openings, to be the key aspect of T cell circumnavigation. Therefore, probing the microenvironment through stochastic, exploratory protrusions into neighboring spaces (chambers) should serve the purpose of mechanosensing the prospects for forming new cortical rings along the migration pass rather than finding the largest space to be in.

### Cell experiments

Primary human CD4+ T cells were isolated from commercially available whole human blood (STEMCELL Technologies Inc., USA, catalog no. 70507.1) with EasySep Human CD4+ T Cell Isolation Kits (STEMCELL Technologies Inc., USA, catalog no. 17952), activated and expanded in ImmunoCult-XF T cell expansion medium (STEMCELL Technologies Inc., USA, catalog no. 10981) with the addition of ImmunoCult human CD3/CD28/CD2 T cell activator and human recombinant interleukin-2 (IL-2; STEMCELL Technologies Inc., USA), as per STEMCELL Technologies Inc. commercial protocol, at 37°C in 5% CO2.

### Polymerization of prestained collagen gels

To prepare collagen-I gel (3 mg/ml), we assembled a gel premix on ice in a prechilled Eppendorf tube. To 1 volume of CellAdhere type I bovine (6 mg/ml, STEMCELL Technologies Inc., USA), we added 8/10 volume of DMEM, 1/10 volume of 10× phosphate-buffered saline (PBS), 1/20 volume of 1 M Hepes, and 1/20 volume of 1 M Atto 647 N-hydroxysuccinimide ester (Sigma-Aldrich, catalog no. 07376), Alexa Fluor 568 ester (Molecular Probes, catalog no. A20003), or Alexa Fluor 488 ester (Molecular Probes, catalog no. A20000). A drop of premixed gel (∼50 μl) was spread immediately on the glass surface of a plasma-treated glass-bottom 35-mm petri dish (MatTek Corp., catalog no. P35G-1.5-14-C) with a pipette tip. During polymerization (room temperature, overnight), gels were covered with 1 ml of mineral oil (Sigma-Aldrich, catalog no. M8410) to prevent evaporation of water. Before adding T cells, polymerized gels were rinsed with PBS to remove the unpolymerized gel components.

### Super-resolution microscopy

For imaging, fixed cells were stained as indicated in the corresponding figures. Imaging was performed using an Instant structured illumination microscopy (iSIM) system (VisiTech Intl, Sunderland, UK) equipped with an Olympus UPLAPO-HR ×100/1.5 NA objective. Image acquisition and system control were done using VisiView® Software (Visitron Systems GmbH, Germany). Images were deconvolved with an iSIM-specific commercial plugin from Microvolution (Cupertino, CA) in FIJI. Figures were composed with Adobe Illustrator CC 2021 (Adobe).

We fixed cells with 4% paraformaldehyde (PFA; Sigma-Aldrich, catalog no. P6148) in1× PBS (Thermo Fisher Scientific, catalog no. 10010023) for the duration of 30 min at room temperature. PFA-fixed cells were then rinsed with 1% bovine serum albumin (BSA) (Thermo Fisher Scientific, catalog no. BP9704) in PBS, followed by 60-minute-long blocking with 1% BSA - 0.5% Triton X-100 (Sigma, catalog no. X100) in PBS. For immunofluorescence staining, all primary antibodies were diluted in 1% BSA PBS. The duration of the incubation with any of the listed primary antibody solutions was 2 hours at room temperature. Similarly, labelings with Alexa Fluor–conjugated secondary antibodies were performed at their final concentration of 5 μg/ml for the duration of 1 hour in 1% BSA PBS at room temperature. After washing out the excess secondary antibodies, chromatin and actin, if necessary, were labeled with 1:1000 Hoechst solution (Tocris, catalog no.5117) and phalloidin conjugates (Sigma-Aldrich, catalog no. 49409; Thermo Fisher Scientific, catalog no. A12380), respectively. We mounted samples using 90% glycerol (Sigma-Aldrich, catalog no. G5516) in 1× PBS.

Antibodies were Phospho-Myosin Light Chain 2 (Ser19) (Cell Signaling, catalog no. 3671), Goat-anti Rabbit IgG Alexa Fluor 488 (Jackson Immuno, catalog no. 111-545-144), and Goat-anti Rabbit IgG Alexa Fluor 568 (Invitrogen, catalog no. A-11036).

## Acknowledgments

We thank C. Combs and D. Malide for the Light Microscopy Core support at the National Heart, Lung, and Blood Institute, NIH. A.S.Z. was supported by the FDA Intramural Research Program of the Center for Biologics Evaluation and Research. A.X.C.-R. acknowledge support from the intramural funding of the Division of Intramural Research Program at the National Institute of Biomedical Imaging and Bioengineering with grant ZIA-EB000094 and the NIH central funds for the NIH Distinguished Scholars Program award. DT was supported by NIH R01 GM136892 and by NSF grant number CMMI 1942561. S. A., D.M.R. and D.V. were supported by NIH grant R35GM136372.

## Supplementary information

**Supplementary Figure 1.**
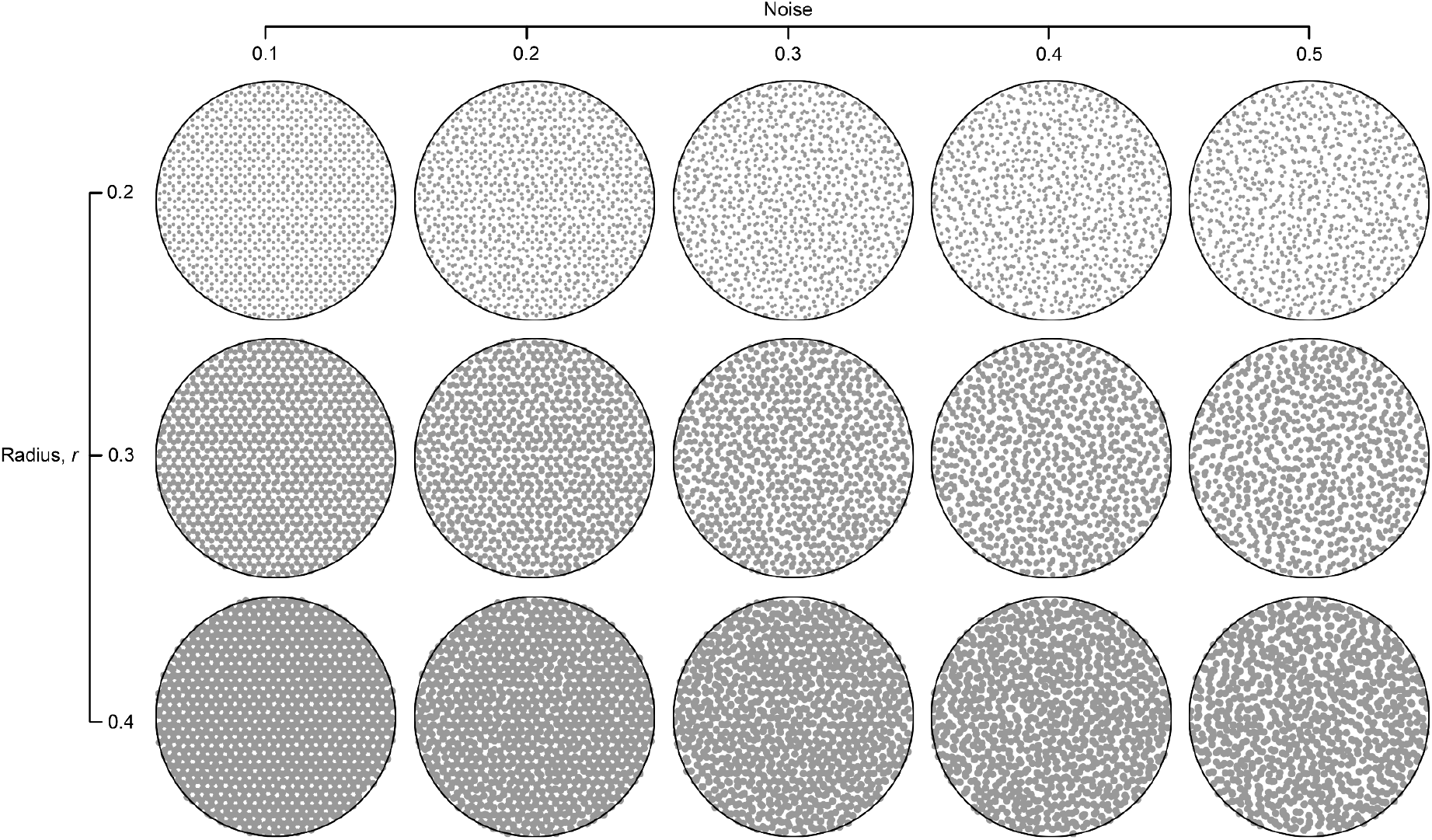
Obstacle field configurations used for T cell migration simulations across varying obstacle radii and spatial noise levels. Circular domains represent two-dimensional obstacle fields with uniformly distributed circular obstacles. Rows correspond to increasing obstacle radii (*r* = 0.2, 0.3, and 0.4, *top to bottom*), while columns correspond to increasing spatial noise levels (0.1 to 0.5, *left to right*). Low noise values result in more ordered, grid-like arrangements, while higher noise values introduce spatial randomness, disrupting uniform spacing and increasing environmental heterogeneity. These synthetic microenvironments were used to systematically evaluate how geometric confinement and obstacle irregularity influence cell motility dynamics.

**Supplementary Table 1.** Simulation Reference Parameter Values (Model of Figures 1-4)

| Parameter | Value | Description |
| --- | --- | --- |
| $\Delta t$ | $3.08 \times 10^{-3} \text{ s}$ | Simulation timestep |
| $L^{Front}$ | $3.60 \text{ }\mu\text{m}$ | Cell leading edge length |
| $\omega$ | $1.00 \times 10^{-3} \text{ s}^{-1}$ | Frequency of fluctuating leading edge bond segment force |
| $l_o$ | $0.270 \text{ }\mu\text{m}$ | Equilibrium bond length |
| $\zeta/l_o$ | $6.38 \times 10^4 \text{ pN}\cdot\text{min}/\mu\text{m}^2$ | Friction coefficient per equilibrium bond length |
| $k^{Area,n}$ | $3 \times 10^5 \text{ pN}/\mu\text{m}$ | Nuclear bond segment spring constant |
| $k^{Area,c}$ | $6000 \text{ pN}/\mu\text{m}$ | Cell membrane bond segment spring constant |
| $k^{Stretch}$ | $1 \times 10^5 \text{ pN}/\mu\text{m}$ | Bond segment spring constant |
| $k^{Excl}$ | $1.69 \times 10^5 \text{ pN}/\mu\text{m}$ | Excluded volume interaction constant |
| $d^{Excl}$ | $0.260 \text{ }\mu\text{m}$ | Repulsive interaction distance between bond segments |
| $F^C$ | $180 - 750 \text{ pN}$ | Magnitude of contractile force range |
| $\kappa$ | $1.53 \text{ pN}\cdot\mu\text{m}^2$ | Bending rigidity |

## Movie legends

**Movie 1** - Comparison of simulated 2D migration of T cells through obstacles of narrow spacing in the absence (*left*) and in the presence of a hydraulic seal around the nucleus. Simulations use the reference parameters listed in Table 1.

**Movie 2** - Comparison of simulated 2D migration of T cells with (*top*) and without (*bottom*) the formation of a hydraulic nuclear piston seal through a field of obstacles of increasing density that is proportional to distance from the entry point. See Table 1 for reference parameters.

**Movie 3** - Comparison of simulated 2D migration of T cells within a randomized obstacle field without (*left*) and with (*right*) the formation of a hydraulic nuclear piston seal, “narrow repolarization” case. Reference parameters are provided in Table 1.

**Movie 4** - Comparison of simulated 2D migration of T cells within a randomized obstacle field without (*left*) and with (*right*) the formation of a hydraulic nuclear piston seal, “wide repolarization” case. Reference parameters are provided in Table 1.

**Movie 5** - Comparison of simulated 2D migration of T cells within a randomized obstacle field without (*left*) and with (*right*) the formation of a hydraulic nuclear piston seal, “no repolarization” case. Reference parameters are provided in Table 1.

**Movie 6** - **A simulation of T cell migration in a confining environment using a model with uropod and high-commitment strategy**.

Here, the T cell is represented with a 3-point model that prohibits direction reversal (due to the presence of the uropod). The large value of the commitment parameter *C* = 20 increases the likelihood that the cell will transmit the nucleus to a new space chamber earlier during the exploratory probing of the environment by the leading protrusion (see the detailed model description in the **Methods** section). Here, the randomized environment is generated with obstacles of radius 0.3 units and their positions perturbed from the hexagonal grid (of unit spacing) by noise of 0.3 units.

**Movie 7** - **A simulation of T cell migration in a confining environment using a model without uropod and high-commitment strategy**

Here, the T cell is represented with a 2-point model that allows for direction reversal (due to the absence of the uropod). The large value of the commitment parameter *C* = 20 increases the likelihood that the cell will transmit the nucleus to a new space chamber earlier during the exploratory probing of the environment by the leading protrusion (see the detailed model description in the **Methods** section). Here, the randomized environment is generated with obstacles of radius 0.3 units and their positions perturbed from the hexagonal grid (of unit spacing) by noise of 0.3 units.

**Movie 8** - **A simulation of T cell migration in a confining environment using a model with uropod and low-commitment strategy**

Here, the T cell is represented with a 3-point model that prohibits direction reversal (due to the presence of the uropod). The small value of the commitment parameter *C* = 1 decreases the likelihood that the cell will transmit the nucleus to a new space chamber until the leading protrusion probes all neighboring chambers (see the detailed model description in the **Methods** section). Here, the randomized environment is generated with obstacles of radius 0.3 units and their positions perturbed from the hexagonal grid (of unit spacing) by noise of 0.3 units.

**Movie 9** - **A simulation of T cell migration in a confining environment using a model without uropod and low-commitment strategy**

Here, the T cell is represented with a 2-point model that allows for direction reversal (due to the absence of the uropod). The small value of the commitment parameter *C* = 1 decreases the likelihood that the cell will transmit the nucleus to a new space chamber until the leading protrusion probes all neighboring chambers (see the detailed model description in the **Methods** section). Here, the randomized environment is generated with obstacles of radius 0.3 units and their positions perturbed from the hexagonal grid (of unit spacing) by noise of 0.3 units.

## References

1. Lämmermann, T. et al. Rapid leukocyte migration by integrin-independent flowing and squeezing. Nature 453, 51–55 (2008).

2. Tabdanov, E. D. et al. Engineering T cells to enhance 3D migration through structurally and mechanically complex tumor microenvironments. Nat. Commun. 12, 2815 (2021).

3. Lämmermann, T. & Sixt, M. Mechanical modes of ‘amoeboid’ cell migration. Curr. Opin. Cell Biol. 21, 636–644 (2009).

4. Marcadis, A. R. et al. Rapid cancer cell perineural invasion utilizes amoeboid migration. Proc. Natl. Acad. Sci. U. S. A. 120, e2210735120 (2023).

5. Liu, Y.-J. et al. Confinement and low adhesion induce fast amoeboid migration of slow mesenchymal cells. Cell 160, 659–672 (2015).

6. Lehmann, S. et al. Hypoxia induces a HIF-1-dependent transition from collective-to-amoeboid dissemination in epithelial cancer cells. Curr. Biol. 27, 392–400 (2017).

7. Te Boekhorst, V. et al. Calpain-2 regulates hypoxia/HIF-induced plasticity toward amoeboid cancer cell migration and metastasis. Curr. Biol. 32, 412–427.e8 (2022).

8. Parlani, M., Jorgez, C. & Friedl, P. Plasticity of cancer invasion and energy metabolism. Trends Cell Biol. 33, 388–402 (2023).

9. Caillier, A., Oleksyn, D., Fowell, D. J., Miller, J. & Oakes, P. W. T cells use focal adhesions to pull themselves through confined environments. J. Cell Biol. 223, (2024).

10. Wolf, K., Müller, R., Borgmann, S., Bröcker, E.-B. & Friedl, P. Amoeboid shape change and contact guidance: T-lymphocyte crawling through fibrillar collagen is independent of matrix remodeling by MMPs and other proteases. Blood 102, 3262–3269 (2003).

11. Ullo, M. F., D’Amico, A. E., Lavenus, S. B. & Logue, J. S. The amoeboid migration of monocytes in confining channels requires the local remodeling of the cortical actin cytoskeleton by cofilin-1. bioRxiv (2023) doi:10.1101/2023.08.11.553020.

12. Reversat, A. et al. Cellular locomotion using environmental topography. Nature 582, 582–585 (2020).

13. Wolf, K. & Friedl, P. Extracellular matrix determinants of proteolytic and non-proteolytic cell migration. Trends Cell Biol. 21, 736–744 (2011).

14. George, S., Martin, J. A. J., Graziani, V. & Sanz-Moreno, V. Amoeboid migration in health and disease: Immune responses versus cancer dissemination. Front. Cell Dev. Biol. 10, 1091801 (2022).

15. Behrooz, A. B. & Shojaei, S. Mechanistic insights into mesenchymal-amoeboid transition as an intelligent cellular adaptation in cancer metastasis and resistance. Biochim. Biophys. Acta Mol. Basis Dis. 1870, 167332 (2024).

16. Friedl, P. & Wolf, K. Plasticity of cell migration: a multiscale tuning model. J. Cell Biol. 188, 11–19 (2010).

17. Graziani, V., Rodriguez-Hernandez, I., Maiques, O. & Sanz-Moreno, V. The amoeboid state as part of the epithelial-to-mesenchymal transition programme. Trends Cell Biol. 32, 228–242 (2022).

18. Yamada, K. M. & Sixt, M. Mechanisms of 3D cell migration. Nat. Rev. Mol. Cell Biol. 20, 738–752 (2019).

19. Driscoll, M. K. et al. Proteolysis-free amoeboid migration of melanoma cells through crowded environments via bleb-driven worrying. Dev. Cell 59, 2414–2428.e8 (2024).

20. Zhovmer, A. S. et al. Septins provide microenvironment sensing and cortical actomyosin partitioning in motile amoeboid T lymphocytes. Sci Adv 10, eadi1788 (2024).

21. Stroka, K. M. et al. Water permeation drives tumor cell migration in confined microenvironments. Cell 157, 611–623 (2014).

22. Aoun, L. et al. Amoeboid Swimming Is Propelled by Molecular Paddling in Lymphocytes. Biophys. J. 119, 1157–1177 (2020).

23. Bergert, M. et al. Force transmission during adhesion-independent migration. Nat. Cell Biol. 17, 524–529 (2015).

24. Charras, G. & Paluch, E. Blebs lead the way: how to migrate without lamellipodia. Nat. Rev. Mol. Cell Biol. 9, 730–736 (2008).

25. Paluch, E., Piel, M., Prost, J., Bornens, M. & Sykes, C. Cortical Actomyosin Breakage Triggers Shape Oscillations in Cells and Cell Fragments. Biophys. J. 89, 724–733 (2005).

26. García-Arcos, J. M. et al. Rigidity percolation and active advection synergize in the actomyosin cortex to drive amoeboid cell motility. Dev. Cell 59, 2990–3007.e7 (2024).

27. Karling, T. & Weavers, H. Immune cells adapt to confined environments in vivo to optimise nuclear plasticity for migration. EMBO Rep. 26, 1238–1268 (2025).

28. Villella, C., Ciccioli, M., Anton, I. M. & Calle, Y. Plasticity in leukocyte migration during haematopoiesis and inflammation. J. Muscle Res. Cell Motil. (2025) doi:10.1007/s10974-025-09691-1.

29. Krummel, M. F., Bartumeus, F. & Gérard, A. T cell migration, search strategies and mechanisms. Nat. Rev. Immunol. 16, 193–201 (2016).

30. Krummel, M. F., Friedman, R. S. & Jacobelli, J. Modes and mechanisms of T cell motility: roles for confinement and Myosin-IIA. Curr. Opin. Cell Biol. 30, 9–16 (2014).

31. Friedl, P. & Weigelin, B. Interstitial leukocyte migration and immune function. Nat. Immunol. 9, 960–969 (2008).

32. Renkawitz, J. et al. Nuclear positioning facilitates amoeboid migration along the path of least resistance. Nature 568, 546–550 (2019).

33. Renkawitz, J. et al. Adaptive force transmission in amoeboid cell migration. Nat. Cell Biol. 11, 1438–1443 (2009).

34. Tooley, A. J. et al. Amoeboid T lymphocytes require the septin cytoskeleton for cortical integrity and persistent motility. Nat. Cell Biol. 11, 17–26 (2008).

35. Adams, G., Jr et al. Functional Behavior of Melanoma Cells Under High Confinement, High Contractility and Low Adhesion Mediating Leader Bleb Based Motility. in MOLECULAR BIOLOGY OF THE CELL vol. 29 (AMER SOC CELL BIOLOGY 8120 WOODMONT AVE, STE 750, BETHESDA, MD 20814-2755 USA, 2018).

36. De Vries, I. J. M. et al. Effective migration of antigen-pulsed dendritic cells to lymph nodes in melanoma patients is determined by their maturation state. Cancer Res. 63, 12–17 (2003).

37. Alvarez, D., Vollmann, E. H. & von Andrian, U. H. Mechanisms and consequences of dendritic cell migration. Immunity 29, 325–342 (2008).

38. Heuzé, M. L. et al. Migration of dendritic cells: physical principles, molecular mechanisms, and functional implications. Immunol. Rev. 256, 240–254 (2013).

39. Ricart, B. G., Yang, M. T., Hunter, C. A., Chen, C. S. & Hammer, D. A. Measuring traction forces of motile dendritic cells on micropost arrays. Biophys. J. 101, 2620–2628 (2011).

40. Lämmermann, T. & Sixt, M. Mechanical modes of ‘amoeboid’cell migration. Curr. Opin. Cell Biol. 21, 636–644 (2009).

41. Sobocinski, G. P. et al. Ultrastructural localization of extracellular matrix proteins of the lymph node cortex: evidence supporting the reticular network as a pathway for lymphocyte migration. BMC Immunol. 11, 42 (2010).

42. Dahl, K. N., Ribeiro, A. J. S. & Lammerding, J. Nuclear shape, mechanics, and mechanotransduction. Circ. Res. 102, 1307–1318 (2008).

43. Rowat, A. C. et al. Nuclear envelope composition determines the ability of neutrophil-type cells to passage through micron-scale constrictions. J. Biol. Chem. 288, 8610–8618 (2013).

44. Denais, C. M. et al. Nuclear envelope rupture and repair during cancer cell migration. Science 352, 353–358 (2016).

45. Friedl, P., Wolf, K. & Lammerding, J. Nuclear mechanics during cell migration. Curr. Opin. Cell Biol. 23, 55–64 (2011).

46. Sánchez-Madrid, F. & Serrador, J. M. Bringing up the rear: defining the roles of the uropod. Nat. Rev. Mol. Cell Biol. 10, 353–359 (2009).

47. Valignat, M.-P. et al. Lymphocytes can self-steer passively with wind vane uropods. Nat. Commun. 5, 5213 (2014).

48. Davidson, P. M., Denais, C., Bakshi, M. C. & Lammerding, J. Nuclear deformability constitutes a rate-limiting step during cell migration in 3-D environments. Cell. Mol. Bioeng. 7, 293–306 (2014).

49. Petrie, R. J., Harlin, H. M., Korsak, L. I. T. & Yamada, K. M. Activating the nuclear piston mechanism of 3D migration in tumor cells. J. Cell Biol. 216, 93–100 (2017).

50. Lee, H.-P. et al. The nuclear piston activates mechanosensitive ion channels to generate cell migration paths in confining microenvironments. Sci. Adv. 7, eabd4058 (2021).

51. Lele, T. P., Dickinson, R. B. & Gundersen, G. G. Mechanical principles of nuclear shaping and positioning. J. Cell Biol. 217, 3330–3342 (2018).

52. Petrie, R. J. & Yamada, K. M. Fibroblasts lead the way: A unified view of 3D cell motility. Trends Cell Biol. 25, 666–674 (2015).

53. Ju, R. J. et al. Compression-dependent microtubule reinforcement enables cells to navigate confined environments. Nat. Cell Biol. 26, 1520–1534 (2024).

54. Lomakin, A. J. et al. The nucleus acts as a ruler tailoring cell responses to spatial constraints. Science 370, eaba2894 (2020).

55. Venturini, V. et al. The nucleus measures shape changes for cellular proprioception to control dynamic cell behavior. Science 370, eaba2644 (2020).

56. Wolf, K. et al. Physical limits of cell migration: control by ECM space and nuclear deformation and tuning by proteolysis and traction force. J. Cell Biol. 201, 1069–1084 (2013).

57. Zhang, X. et al. SUN1/2 and Syne/Nesprin-1/2 complexes connect centrosome to the nucleus during neurogenesis and neuronal migration in mice. Neuron 64, 173–187 (2009).

58. Lombardi, M. L. et al. The interaction between nesprins and sun proteins at the nuclear envelope is critical for force transmission between the nucleus and cytoskeleton. J. Biol. Chem. 286, 26743–26753 (2011).

59. Thiam, H.-R. et al. Perinuclear Arp2/3-driven actin polymerization enables nuclear deformation to facilitate cell migration through complex environments. Nat. Commun. 7, 10997 (2016).

60. Hu, D. J.-K. et al. Dynein recruitment to nuclear pores activates apical nuclear migration and mitotic entry in brain progenitor cells. Cell 154, 1300–1313 (2013).

61. Tsai, M.-H. et al. Impairment in dynein-mediated nuclear translocation by BICD2 C-terminal truncation leads to neuronal migration defect and human brain malformation. Acta Neuropathol Commun 8, 106 (2020).

62. Baffet, A. D., Hu, D. J. & Vallee, R. B. Cdk1 activates pre-mitotic nuclear envelope dynein recruitment and apical nuclear migration in neural stem cells. Dev. Cell 33, 703–716 (2015).

63. Marks, P. C., Hewitt, B. R., Baird, M. A., Wiche, G. & Petrie, R. J. Plectin linkages are mechanosensitive and required for the nuclear piston mechanism of three-dimensional cell migration. Mol. Biol. Cell 33, ar104 (2022).

64. Petrie, R. J., Koo, H. & Yamada, K. M. Generation of compartmentalized pressure by a nuclear piston governs cell motility in a 3D matrix. Science 345, 1062–1065 (2014).

65. Tsai, J.-W., Lian, W.-N., Kemal, S., Kriegstein, A. R. & Vallee, R. B. Kinesin 3 and cytoplasmic dynein mediate interkinetic nuclear migration in neural stem cells. Nat. Neurosci. 13, 1463–1471 (2010).

66. Friedl, P., Entschladen, F., Conrad, C., Niggemann, B. & Zänker, K. S. CD4+ T lymphocytes migrating in three-dimensional collagen lattices lack focal adhesions and utilize beta1 integrin-independent strategies for polarization, interaction with collagen fibers and locomotion. Eur. J. Immunol. 28, 2331–2343 (1998).

67. Caillier, A., Oleksyn, D., Fowell, D. J., Miller, J. & Oakes, P. W. T cells use focal adhesions to pull themselves through confined environments. bioRxivorg (2023) doi:10.1101/2023.10.16.562587.

68. Chabaud, M. et al. Cell migration and antigen capture are antagonistic processes coupled by myosin II in dendritic cells. Nat. Commun. 6, 7526 (2015).

69. Vargas, P. et al. Innate control of actin nucleation determines two distinct migration behaviours in dendritic cells. Nat. Cell Biol. 18, 43–53 (2016).

70. Wong, K., Van Keymeulen, A. & Bourne, H. R. PDZRhoGEF and myosin II localize RhoA activity to the back of polarizing neutrophil-like cells. J. Cell Biol. 179, 1141–1148 (2007).

71. Morin, N. A. et al. Nonmuscle myosin heavy chain IIA mediates integrin LFA-1 de-adhesion during T lymphocyte migration. J. Exp. Med. 205, 195–205 (2008).

72. Smith, A., Bracke, M., Leitinger, B., Porter, J. C. & Hogg, N. LFA-1-induced T cell migration on ICAM-1 involves regulation of MLCK-mediated attachment and ROCK-dependent detachment. J. Cell Sci. 116, 3123–3133 (2003).

73. Smith, L. A., Aranda-Espinoza, H., Haun, J. B., Dembo, M. & Hammer, D. A. Neutrophil traction stresses are concentrated in the uropod during migration. Biophys. J. 92, L58–60 (2007).

74. Uchida, K. S. K., Kitanishi-Yumura, T. & Yumura, S. Myosin II contributes to the posterior contraction and the anterior extension during the retraction phase in migrating Dictyostelium cells. J. Cell Sci. 116, 51–60 (2003).

75. Rochussen, A. M. et al. Actin conformation dynamics precede force generation in cytotoxic T lymphocytes. bioRxiv 2025.05.15.654337 (2025) doi:10.1101/2025.05.15.654337.

76. Ratner, S., Sherrod, W. S. & Lichlyter, D. Microtubule retraction into the uropod and its role in T cell polarization and motility. J. Immunol. 159, 1063–1067 (1997).

77. Etienne-Manneville, S. Microtubules in Cell Migration. Annu. Rev. Cell Dev. Biol. 29, 471–499 (2013).

78. Hind, L. E., Vincent, W. J. B. & Huttenlocher, A. Leading from the back: The role of the uropod in neutrophil polarization and migration. Dev. Cell 38, 161–169 (2016).

79. Xu, J. et al. Divergent signals and cytoskeletal assemblies regulate self-organizing polarity in neutrophils. Cell 114, 201–214 (2003).

80. Caudron, F. & Barral, Y. Septins and the lateral compartmentalization of eukaryotic membranes. Dev. Cell 16, 493–506 (2009).

81. Hu, Q. et al. A septin diffusion barrier at the base of the primary cilium maintains ciliary membrane protein distribution. Science 329, 436–439 (2010).

82. Pouwels, J. et al. SHARPIN regulates uropod detachment in migrating lymphocytes. Cell Rep. 5, 619–628 (2013).

83. Alam, S. G. et al. The nucleus is an intracellular propagator of tensile forces in NIH 3T3 fibroblasts. J. Cell Sci. 128, 1901–1911 (2015).

84. DiNapoli, K. T., Robinson, D. N. & Iglesias, P. A. A mesoscale mechanical model of cellular interactions. Biophys. J. 120, 4905–4917 (2021).

85. Echevarría, W., Leite, M. F., Guerra, M. T., Zipfel, W. R. & Nathanson, M. H. Regulation of calcium signals in the nucleus by a nucleoplasmic reticulum. Nat. Cell Biol. 5, 440–446 (2003).

86. Galva, C., Artigas, P. & Gatto, C. Nuclear Na+/K+-ATPase plays an active role in nucleoplasmic Ca2+ homeostasis. J. Cell Sci. 125, 6137–6147 (2012).

87. Kiessling, M., Djalinac, N., Voglhuber, J. & Ljubojevic-Holzer, S. Nuclear calcium in cardiac (patho)physiology: Small compartment, big impact. Biomedicines 11, 960 (2023).

88. Resende, R. R. et al. Nucleoplasmic calcium signaling and cell proliferation: calcium signaling in the nucleus. Cell Commun. Signal. 11, 14 (2013).

89. Badalamenti, G. et al. Role of tumor-infiltrating lymphocytes in patients with solid tumors: Can a drop dig a stone? Cell. Immunol. 343, 103753 (2019).

90. Chai, L. F., Prince, E., Pillarisetty, V. G. & Katz, S. C. Challenges in assessing solid tumor responses to immunotherapy. Cancer Gene Ther. 27, 528–538 (2020).

91. Kuczek, D. E. et al. Collagen density regulates the activity of tumor-infiltrating T cells. J Immunother Cancer 7, 68 (2019).

92. Zhao, R. et al. Targeting the Microtubule-Network Rescues CTL Killing Efficiency in Dense 3D Matrices. Front. Immunol. 12, 729820 (2021).

93. Song, J. et al. The extracellular matrix of lymph node reticular fibers modulates follicle border interactions and germinal center formation. iScience 26, 106753 (2023).

94. Sadjadi, Z., Zhao, R., Hoth, M., Qu, B. & Rieger, H. Migration of Cytotoxic T Lymphocytes in 3D Collagen Matrices. Biophys. J. 119, 2141–2152 (2020).

95. Pruitt, H. C. et al. Collagen fiber structure guides 3D motility of cytotoxic T lymphocytes. Matrix Biol. 85-86, 147–159 (2020).

96. Salmon, H. et al. Matrix architecture defines the preferential localization and migration of T cells into the stroma of human lung tumors. J. Clin. Invest. 122, 899–910 (2012).

97. Binnewies, M. et al. Understanding the tumor immune microenvironment (TIME) for effective therapy. Nat. Med. 24, 541–550 (2018).

98. Joyce, J. A. & Fearon, D. T. T cell exclusion, immune privilege, and the tumor microenvironment. Science 348, 74–80 (2015).

99. Guha, P., Heatherton, K. R., O’Connell, K. P., Alexander, I. S. & Katz, S. C. Assessing the future of solid tumor immunotherapy. Biomedicines 10, 655 (2022).

100. Wu, B., Zhang, B., Li, B., Wu, H. & Jiang, M. Cold and hot tumors: from molecular mechanisms to targeted therapy. Signal Transduct. Target. Ther. 9, 274 (2024).

101. Lanitis, E., Dangaj, D., Irving, M. & Coukos, G. Mechanisms regulating T-cell infiltration and activity in solid tumors. Ann. Oncol. 28, xii18–xii32 (2017).

102. Donnadieu, E., Dupré, L., Pinho, L. G. & Cotta-de-Almeida, V. Surmounting the obstacles that impede effective CAR T cell trafficking to solid tumors. J. Leukoc. Biol. 108, 1067–1079 (2020).

103. Slaney, C. Y., Kershaw, M. H. & Darcy, P. K. Trafficking of T cells into tumors. Cancer Res. 74, 7168–7174 (2014).

104. Tagay, Y. et al. Dynein-powered cell locomotion guides metastasis of breast cancer. Adv. Sci. e2302229 (2023).

105. Zhovmer, A. S. et al. Mechanical Counterbalance of Kinesin and Dynein Motors in a Microtubular Network Regulates Cell Mechanics, 3D Architecture, and Mechanosensing. ACS Nano 15, 17528–17548 (2021).

106. Bufi, N. et al. Human primary immune cells exhibit distinct mechanical properties that are modified by inflammation. Biophys. J. 108, 2181–2190 (2015).

107. Rutkowski, D. M. & Vavylonis, D. Discrete mechanical model of lamellipodial actin network implements molecular clutch mechanism and generates arcs and microspikes. PLoS Comput. Biol. 17, e1009506 (2021).

108. Treado, J. D. et al. Bridging particle deformability and collective response in soft solids. Phys. Rev. Mater. 5, (2021).

109. Kim, T., Hwang, W., Lee, H. & Kamm, R. D. Computational analysis of viscoelastic properties of crosslinked actin networks. PLoS Comput. Biol. 5, e1000439 (2009).

110. Stöberl, S. et al. Nuclear deformation and dynamics of migrating cells in 3D confinement reveal adaptation of pulling and pushing forces. bioRxiv 2023.10.30.564765 (2023) doi:10.1101/2023.10.30.564765.

